# Multiple mechanisms impact fluconazole resistance of mutant Erg11 proteins in Candida glabrata

**DOI:** 10.1101/2021.06.23.449691

**Authors:** Bao Gia Vu, W. Scott Moye-Rowley

## Abstract

Azoles, the most commonly used antifungal drugs, specifically inhibit the fungal lanosterol α-14 demethylase enzyme, which is referred to as Erg11. Inhibition of Erg11 ultimately leads to a reduction in ergosterol production, an essential fungal membrane sterol. Many *Candida* species, such as *Candida albicans*, develop mutations in this enzyme which reduces the azole binding affinity and results in increased resistance. *Candida glabrata* is also a pathogenic yeast that has low intrinsic susceptibility to azole drugs and easily develops elevated resistance. These azole resistant mutations typically cause hyperactivity of the Pdr1 transcription factor and rarely lie within the *ERG11* gene. Here, we generated *C. glabrata ERG11* mutations that were analogous to azole resistance alleles from *C. albicans ERG11*. Three different Erg11 forms (Y141H, S410F, and the corresponding double mutant (DM)) conferred azole resistance in *C. glabrata* with the DM Erg11 form causing the strongest phenotype. The DM Erg11 also induced cross-resistance to amphotericin B and caspofungin. Resistance caused by the DM allele of *ERG11* imposed a fitness cost that was not observed with hyperactive *PDR1* alleles. Crucially, the presence of the DM *ERG11* allele was sufficient to activate the Pdr1 transcription factor in the absence of azole drugs. Our data indicate that azole resistance linked to changes in *ERG11* activity can involve cellular effects beyond an alteration in this key azole target enzyme. Understanding the physiology linking ergosterol biosynthesis with Pdr1-mediated regulation of azole resistance is crucial for ensuring the continued efficacy of azole drugs against *C. glabrata*.

**Importance:** Azole drugs target the Erg11 enzyme and lead to a reduction in fungal ergosterol, a vital sterol in yeast. Mutations in Erg11 are common among azole resistant *Candida albicans* clinical isolates, but not in *C. glabrata*, a major human pathogen. In this study, we showed that *ERG11* mutations were tolerated in *C. glabrata*, and these mutations could confer azole resistance. We found that the strongest azole-resistant allele of *ERG11* led to induction of the Pdr1 transcription factor and Cdr1 ATP-binding cassette transporter protein in the absence of drug. *ERG11* mutations can cause azole resistance via altered enzymatic properties but also by triggering induction of other resistance systems owing to impacts on ergosterol biosynthesis. These data illustrate the deep connections between ergosterol biosynthesis and regulation of membrane transporter proteins via Pdr1 and the ergosterol-responsive transcription factor Upc2A.

## Introduction

Ergosterol is essential for fungal plasma membrane homeostasis (1, 2). It maintains membrane integrity and fluidity, as well as facilitates membrane-bound enzymatic reactions (1). The azole class of antifungal therapy targets the fungal ability to synthesize ergosterol through inhibition of the function of lanosterol 14-alpha-demethylase, which is encoded by the *ERG11* gene (3) (4). The Erg11 enzyme catalyzes a three-step reaction that results in the demethylation of lanosterol. In each step, one oxygen molecule is oxidized, leading to the production of one NADPH molecule (4, 5). The nucleophilic N-4 atom of the azole ring binds to heme, while the N-1 group interacts with the enzyme active site, directly competing with the normal substrate (lanosterol) binding (6, 7). While azoles bind selectively to the Erg11 enzyme (8) the affinity of this enzyme for different azoles varies; with fluconazole binding with lower affinity than either itraconazole or voriconazole (4).

In *Candida albicans* (Ca), mutations in the Erg11 enzyme have been shown to alter azole binding capacity and significantly reduce the drug inhibitory effect resulting in an enhanced azole resistant phenotype (9–14)}. The majority of these mutations cluster into three hot spots, located within residues 105 to 165, 266 to 287, and 405 to 488 (15). The ability to confer azole resistance by some of these mutations has also been confirmed by introducing these mutations on plasmids into a heterologous *Saccharomyces cerevisiae* system and then assessing azole susceptibility. With this approach, single amino acid mutations, such as: Y132H, S405F, G464S, R467K, and double mutations, such as: Y132H with S405F, Y132H with G464S, R467K with G464S have been shown to confer azole resistance in *S. cerevisiae* (9). In addition, several *CaERG11* constructs of the enzyme catalytic domain containing amino acid substitutions, including Y132H, F145L, I471T, S279F, and G464S have been expressed in *Escherichia coli*, purified, and showed to exhibit a lower binding capacity to azoles (15). Finally, many of these mutant alleles have also been introduced back into an azole-susceptible *C. albicans* strain, at the *ERG11* gene native locus, to confirm the relative contributions of individual and combined mutation to azole resistance (10).

Unlike in *C. albicans*, mutations in the *ERG11* gene have been rarely reported among *Candida glabrata* azole resistant clinical isolates. The most common amino acid substitution mutations in *C. glabrata* are in the gene encoding a Zn_2_Cys_6_ zinc cluster-containing transcription factor called Pdr1 (16). These mutations yield a gain-of-function (GOF) phenotype and lead to the elevated transcription of downstream target genes, such as the ATP-binding cassette (ABC) transporter-encoding *CDR1* gene. The Cdr1 transporter is required for the azole resistant mechanism in both wild-type and GOF *PDR1 C. glabrata* isolates (17).

We recently reported that there is biological crosstalk between the ergosterol biosynthesis pathway and the drug efflux pump system in *C. glabrata* (18). Alteration in ergosterol synthesis directly induced expression of both Pdr1 and Cdr1. The Zn_2_Cys_6_ zinc cluster-containing factor Upc2A is essential for such cross-interaction, through its functions as a master transcription factor regulating the expression levels of both ergosterol synthesis genes (*ERG*) and *PDR1*-*CDR1* genes (19) (20). Since our earlier experiments indicated that activity of the Pdr1/*CDR1* system was responsive to ergosterol biosynthesis, we further wanted to determine if mutations in *ERG11* that reduced azole susceptibility might act via induction of this separate azole resistance pathway. Considering the conserved inhibitory effects of azoles on the Erg11 protein in both *C. albicans* and *C. glabrata*, we first generated isogenic *C. glabrata* isolates carrying mutant *ERG11* alleles, analogs of the *CaERG11* Y132H, S405F or Y132H-S405F mutations. Then we examined the effects of these mutant alleles on ergosterol synthesis, antifungal drug resistance, and abilities to cross-regulate the drug efflux pump Pdr1-Cdr1 system. Our data indicate that azole-resistant *ERG11* mutations can activate function of Pdr1 with attendant induction of genes (like *CDR1*) downstream of this factor, even in the absence of drug. These findings illustrate the importance of considering secondary effects of *ERG11* mutations on activation of other resistance pathways when evaluating the azole resistance caused by these lesions. Our data provide further evidence directly tying expression of the Pdr1-dependent resistance pathway to ergosterol biosynthesis.

## Results

### Generation of azole-resistant forms of *C. glabrata* Erg11

Detailed structural information is available for the Erg11 enzymes from these two pathogenic *Candida* species as their three dimensional structures have recently been solved by X-ray crystallography, along with the previously determined structure of *S. cerevisiae* Erg11 (6, 21). Figure 1A shows an alignment of the amino acid sequences of these three closely related proteins.

**Figure 1.**
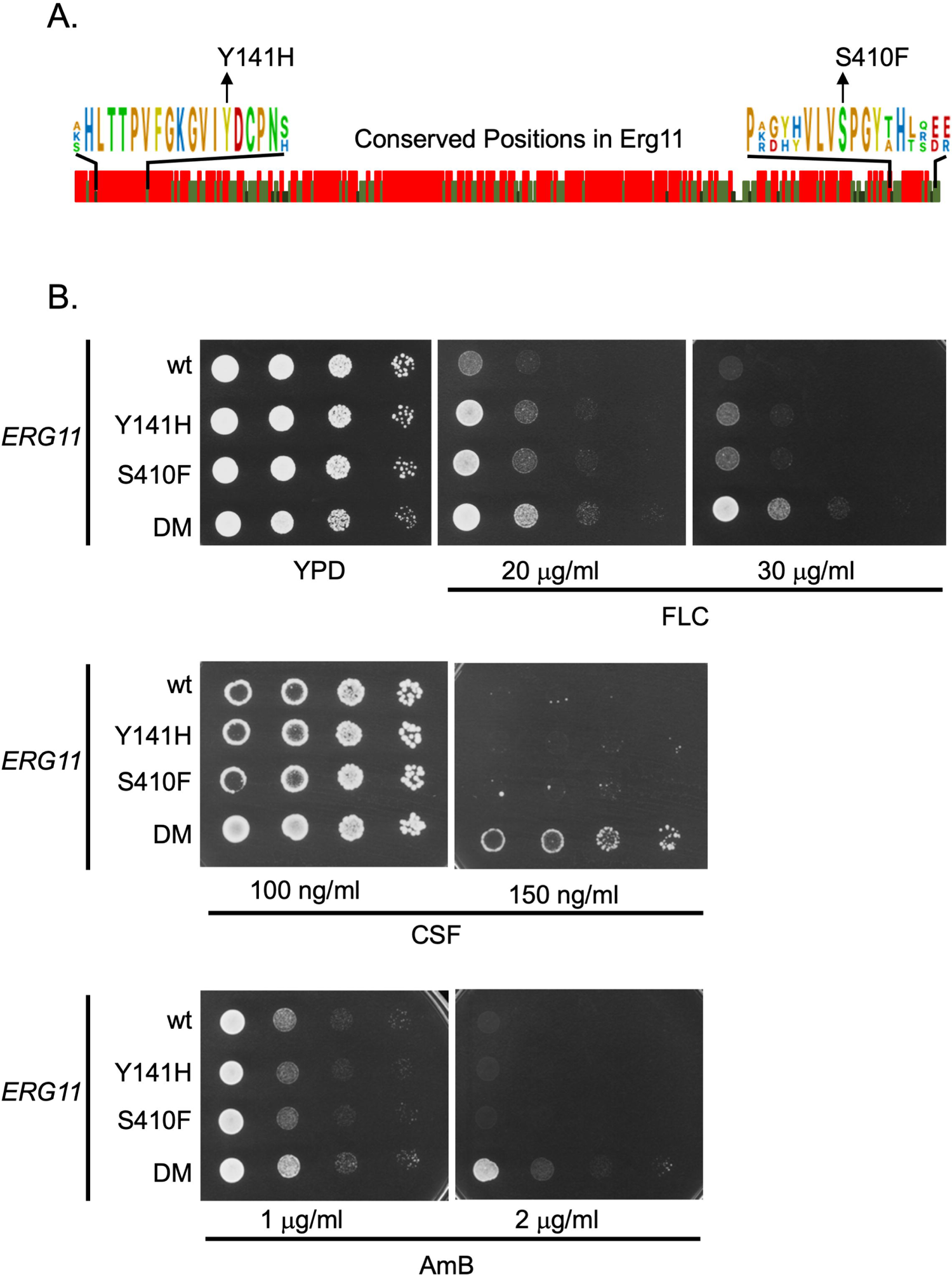
Construction of mutant *ERG11* alleles in *Candida glabrata*. A. Amino acid sequence comparison between Erg11 proteins from *Saccharomyces cerevisiae*, *Candida glabrata* and *Candida albicans*. Graphic indication of amino acid identity between these three yeast enzymes is shown with a red bar indicating 100%, while light and dark green denotes 66% or no conservation. Expanded regions with conserved residues are shown to illustrate the positions that were mutated in *C. glabrata* to match changes associated with fluconazole resistance originally found in *C. albicans*. B. Isogenic *C. glabrata* strains containing the indicated *ERG11* alleles were grown to mid-log phase and plated as serial dilutions on rich medium (YPD) or the same medium containing the indicated concentrations of antifungal drugs. The strain containing both the Y141H- and S410F-encoding alleles was designated double mutant (DM). Plates were allowed to develop at 30°C and then photographed. Antifungal drugs are abbreviated as fluconazole (FLC), caspofungin (CSF) and amphotericin B (AmB).

Previous analyses of azole resistant clinical isolates of *C. albicans* identified changes in residue tyrosine 132 to histidine (Y132H), serine 405 to phenylalanine (S405F) and a double mutant containing both of these lesions (Y132H S405F, referred to as double mutant or DM here) to trigger a decrease in fluconazole susceptibility when present in the *C. albicans* Erg11 enzyme (9). Inspection of the aligned sequences shown in Figure 1A indicated that both Y141 and S410 (in *C. glabrata*) were conserved residues across the enzymes from *S. cerevisiae* and the two *Candida* species. To determine if these same substitution mutations would cause similar fluconazole resistance phenotypes when introduced into the *C. glabrata* Erg11, we used site-directed mutagenesis to generate the appropriate mutant enzymes. These were returned to *C. glabrata* as the sole source of Erg11 using a plasmid shuffling approach (see Materials and Methods). Appropriate transformants were tested for their resistance to fluconazole, caspofungin and amphotericin B (Figure 1B).

The presence of either single mutant form of Erg11 (Y141H or S410F) led to a slight increase in fluconazole resistance compared to the wild-type enzyme while the double mutant (Y141H and S410F: DM) caused the strongest elevation in fluconazole resistance. The enhanced effect on azole resistance caused by combining these two mutations into a single enzyme has been seen before (9). Only the DM Erg11 was able to enhance resistance to caspofungin or amphotericin B. These data indicated that these mutant forms of *C. glabrata* Erg11 were able to increase fluconazole resistance like their *C. albicans* counterparts and, when combined in the DM Erg11, increased resistance to two other antifungal drugs. We also tested these strains for their susceptibility to voriconazole and itraconazole (Supplemental Figure 1) and found that the mutants caused voriconazole phenotypes similar to those seen for fluconazole but had no significant effect on itraconazole susceptibility. This differential response of a given *ERG11* mutant to different azole drugs has been observed previously (9, 22), with relatively large effects on FLC resistance not correlating with a similar effect to itraconazole susceptibility.

The multidrug resistant nature of the DM *ERG11* mutant strain was not expected. This finding prompted us to evaluate the levels of ergosterol supported by each of these mutant strains as well as expression of genes that could explain the increased resistance to caspofungin. The three mutant strains along with their wild-type *ERG11* counterpart were grown to mid-log phase and allowed to grow for 3 or 24 hours with or without fluconazole challenge.

Total sterols were extracted from these cultures and assayed for the level of ergosterol produced in each strain. Total RNA was also prepared from isogenic wild-type and DM *ERG11* strains without fluconazole challenge and assayed by RT-qPCR for the mRNA levels of the genes indicated.

Ergosterol levels were the lowest in the DM Erg11-expressing strains in the absence of fluconazole stress (Figure 2A). The two single mutants produced ergosterol levels that were very similar to those seen in the wild-type strain. A 3 hour treatment with fluconazole caused a dramatic change in relative ergosterol levels as at this point, the DM Erg11 strain had the highest level of ergosterol while the other 3 strains dropped below this level (Figure 2B). No differences were observed between the two mutant forms of Erg11 compared to the wild-type protein until the fluconazole treatment was extended to 24 hours (Figure 2C). At this time, the DM Erg11 still produced the highest level of ergosterol but now both the Y141H and S410F single mutant strains had levels of ergosterol that were higher than the wild-type but still lower than the DM Erg11.

**Figure 2.**
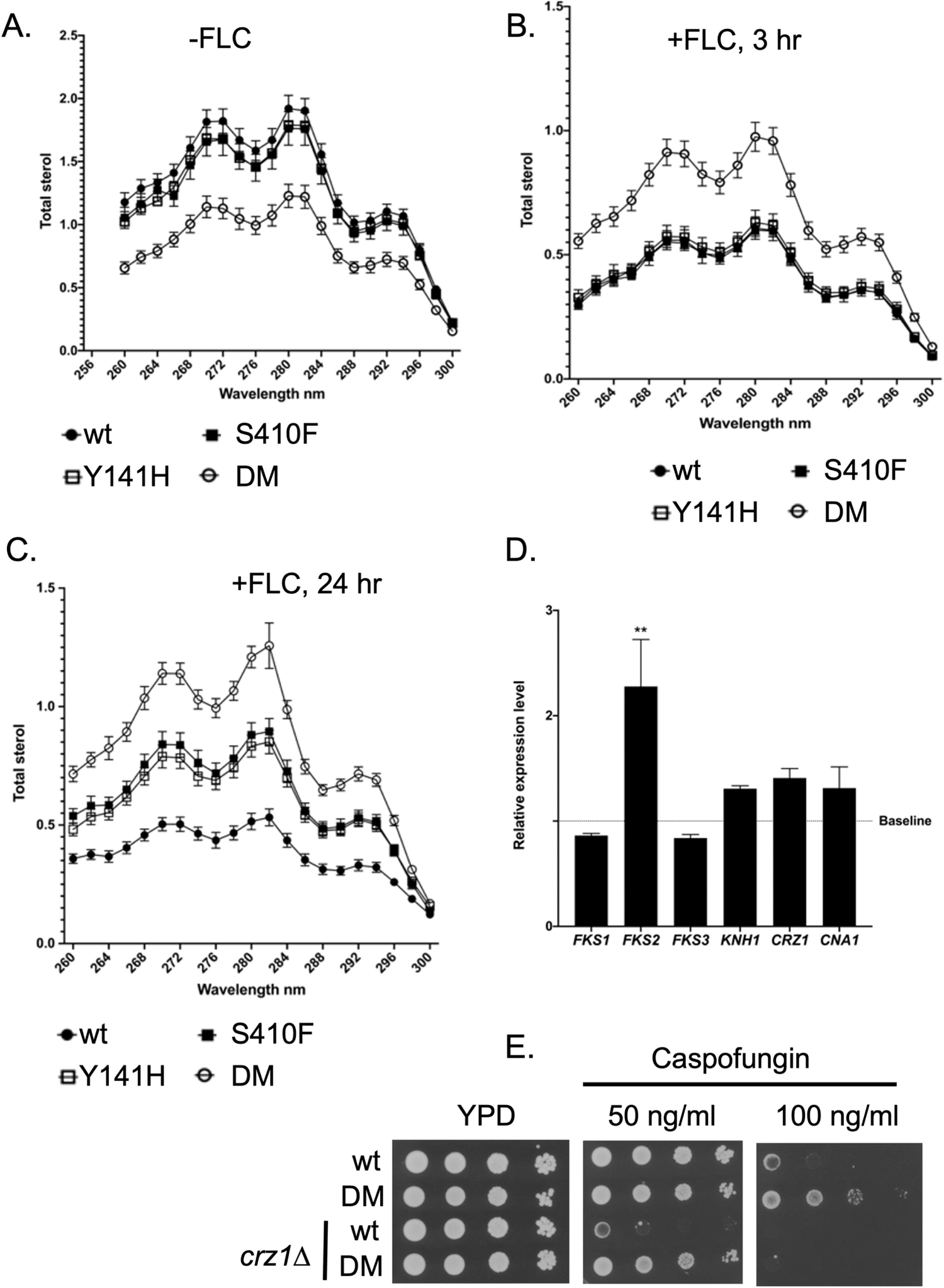
Ergosterol levels and caspofungin resistance-related mRNA production in *ERG11* mutant strains. Mid-log phase *C. glabrata* strains containing the indicated *ERG11* alleles were either not treated (A) with fluconazole (20 μg/ml) (-FLC) or exposed to this drug for 3 (B) or 24 (C) hours. Total lipid extracts were prepared and analyzed for the levels of ergosterol in each sample. D. Total RNA was extracted from isogenic wild-type and DM *ERG11* strains without fluconazole treatment. Samples were then assayed for levels of the indicated mRNAs using RT-qPCR. Relative expression level refers to the ratio of mRNA produced in DM *ERG11* strains/wild-type strain. Baseline indicates equivalent expression in the wildtype strain. E. Loss of *CRZ1* prevents increased caspofungin resistance of a DM *ERG11* strain. The *CRZ1* gene was disrupted in wild-type and DM *ERG11* strains. These *crz1Δ* derivatives and their isogenic wild-type strains were grown to mid-log phase and then analyzed by serial dilution on YPD plates without (YPD) or containing the indicated levels of caspofungin. Plates were incubated at 30°C for two days and photographed.

Analysis of mRNA levels from genes directly involved in production of β-glucan synthase (*FKS1*, *FKS2*, *FKS3*, *KNH1*) as well as regulators of these genes (*CRZ1*, *CNA1*) indicated that only the mRNA for the *FKS2* gene was elevated in the DM *ERG11* mutant compared to its cognate wild-type strain (Figure 2D). This transcriptional induction of *FKS2* could serve to explain the increase in caspofungin resistance seen in the DM Erg11-expressing strain.

We analyzed the contribution of the Crz1 transcription factor to the DM Erg11-triggered increase in caspofungin resistance by disrupting this gene in both wild-type and the isogenic DM *ERG11* strains (Figure 2E). At low concentrations of caspofungin, DM Erg11 was still able to increase drug resistance in a *crz1Δ* strain compared to the wild-type *ERG11* (see 50 ng/ml plate). However, loss of Crz1 eliminated high level (100 ng/ml) caspofungin resistance. We interpret these data to suggest there are at least two effects of the DM *ERG11* allele on caspofungin resistance: a minor one that is Crz1 independent and a major one that requires the presence of Crz1.

### Communication between *ERG11* and Pdr1/*CDR1* pathway

Our previous work demonstrated that reduction in Erg11 expression led to an induction of the Pdr1 transcription factor and its target genes such as *CDR1* (19). To determine if these mutant forms of Erg11 triggered a similar response, we measured expression of Pdr1 and Cdr1 at both the protein and mRNA levels. Transformants containing the various *ERG11* alleles were grown to mid-log phase and either treated with fluconazole for 3 hours or left untreated. Total RNA was prepared from these cultures and mRNA for *CDR1*, *PDR1* and *ERG11* measured by RT-qPCR (Figure 3A).

**Figure 3.**
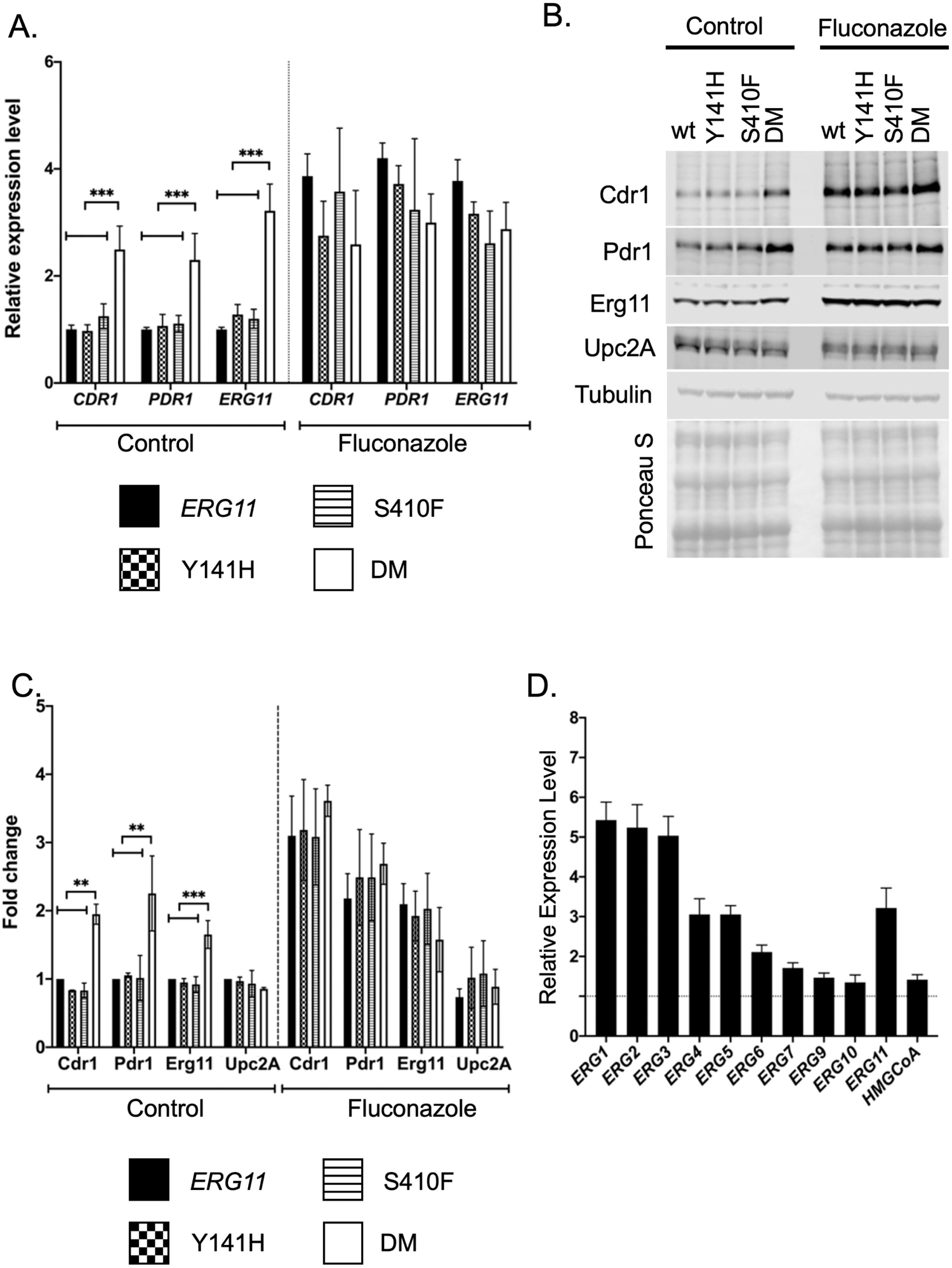
Expression changes in response to *ERG11* alleles. A. Levels of mRNA for *PDR1, CDR1,* and *ERG11* were assayed by RT-qPCR in the indicated *ERG11* mutant strains during growth in the absence (Control) or presence of fluconazole stress (Fluconazole). Fluconazole treatment was 20 μg/ml for 2 hours. B. Whole cell protein extracts were prepared from the same strains in A. These extracts were analyzed by western blotting using the indicated antisera against Cdr1, Pdr1, Erg11, Upc2A or Tubulin. Equal loading and transfer of these extracts was ensured by staining the nitrocellulose membrane with Ponceau S prior to antisera incubation. C. Quantitation of 4 replicates of this western blotting is shown with the data presented as the ratio of each strain relative to the wild-type cells. D. RT-qPCR analysis of *ERG* gene expression in the DM *ERG11* mutant background relative to the wild-type level. HMGCoA (HMG-CoA reductase) refers to the *HMG1* or *CAGL0L11506g* locus.

In the absence of fluconazole stress, only the DM *ERG11* allele led to induction of all three of these mRNAs. Upon challenge with fluconazole, all of these genes were induced to similar relative levels. The enhanced fluconazole resistance seen in the DM *ERG11* strain may be explained by elevated *PDR1* and *CDR1* transcription, at least in part.

The levels of these proteins were also evaluated by western blots using appropriate rabbit polyclonal antibodies for each *C. glabrata* protein (Figure 3B) and quantitated (Figure 3C). Both Pdr1 and Cdr1 were elevated in untreated cells along with Erg11 in the DM *ERG11* strain. Treatment with fluconazole led to similar levels of induction of all these proteins. Upc2A protein levels were not changed under any of these conditions or genetic backgrounds. This is consistent with Upc2A regulation occurring at the post-translational step as has been suggested (23–25).

The levels of mRNA of a range of genes in the ergosterol biosynthetic pathway were also determined in both the DM and wild-type Erg11 strains (Figure 3D). These data are presented as a ratio of mRNA level in the DM Erg11 strain over the wild-type level for each transcript. Most of these *ERG* genes were induced at least 2-fold in the DM Erg11-expressing cells, consistent with the ergosterol limitation in this strain although several *ERG* genes including *ERG9*, *ERG7*, *ERG10* and *HMG1* showed little to no increase. Clearly, the presence of the DM Erg11 enzyme induced a wide range of transcriptional activation across the ergosterol pathway in addition to its effects on *PDR1* and *CDR1* expression. We also suggest that the DM *ERG11* strain may activate Crz1 leading to the previously observed increase in *FKS2* expression and caspofungin resistance. These effects on Pdr1 and Crz1 would explain the multidrug resistance seen in DM *ERG11* mutant strains.

### Upc2A function is increased in the *ERG11* mutants

Our previous data showed that genetic inhibition of *ERG11* gene expression could induce the expression of the drug efflux pump system Pdr1-Cdr1 (19). Upc2A was essential for this interaction in which it directly induced the expression levels of genes in both the *ERG* and Pdr1/*CDR1* pathways (19, 20). Since these two pathways were also induced in the DM *ERG11* mutant, we wanted to determine if Upc2A function was activated in response to these changes in the *ERG11* gene. To test this, we generated an Upc2A reporter construct that contains five copies of the Upc2A binding site (Sterol Response Element: SRE) from the *ERG1* promoter. These concatemerized SREs were placed upstream from the *S. cerevisiae CYC1* promoter that was fused to *E. coli lacZ*. To confirm that this reporter system was sensitive to the *UPC2A* allele, we introduced this plasmid into isogenic *C. glabrata* strains containing the 3X HA*-UPC2A*, 3X HA*-*G898D *UPC2A* gain-of-function mutation (20) or a null allele of *UPC2A*. A plasmid containing only the *CYC1-lacZ* fusion gene with no upstream activation sequence was also used as negative control (Figure 4A).

**Figure 4.**
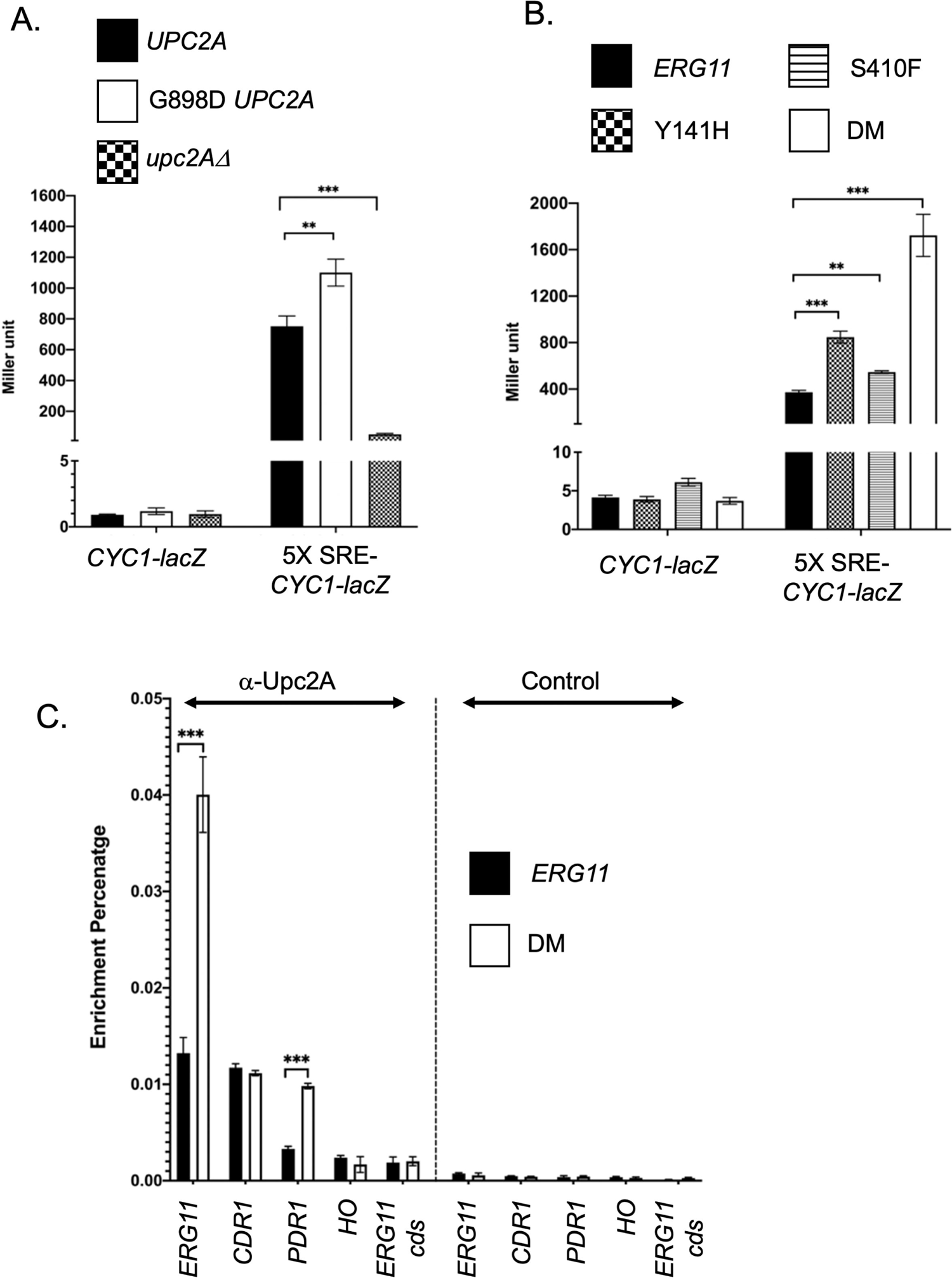
Regulation of Upc2A in response to *ERG11* alleles. A. Two different reporter constructs carried on low-copy-number *C. glabrata* vectors that contained either the *S. cerevisiae CYC1* promoter region with a translational fusion to *E. coli lacZ* (*CYC1-lacZ*) or the same reporter construct with five copies of the *C. glabrata ERG1* sterol response element (SRE) cloned upstream of the *CYC1* promoter (5X SRE-*CYC1-lacZ*) were introduced into the 3 different *C. glabrata* strains indicated at the top. These strains varied at their *UPC2A* allele corresponding to a wild-type strain (*UPC2A*), gain-of-function form of *UPC2A* (G898D *UPC2A*) or a null mutation (*upc2AΔ*). Transformants were grown to mid-log phase and then assayed for the level of β-galactosidase produced in each strain. B. The two different reporter plasmids from A were introduced into the 4 isogenic *ERG11* mutant strains indicated. Transformants were grown to mid-log phase and assayed for β-galactosidase levels. C. Chromatin immunoprecipitation (ChIP) of Upc2A DNA-binding to genomic target sites. Total sheared and crosslinked chromatin was prepared from either wild-type or DM *ERG11* cells. ChIP reactions were carried out with either anti-Upc2A or a non-specific control antisera. Immunopurified DNA was quantitated with qPCR using primer pairs that detected the promoters of *ERG11* (*ERG11*), *CDR1* (*CDR1*), *PDR1* (*PDR1*) or *HO* (*HO*) along with a primer pair that detected a segment of the *ERG11* coding sequence (*ERG11* cds). Data are plotted as the percentage of DNA recovered in the immunopurified sample/total input DNA.

The introduction of the gain-of-function form of Upc2A (G898D *UPC2A*) led to production of the highest level of β-galactosidase seen from the 5X SRE-*CYC1-lacZ* plasmid with nearly 1200 Miller units of activity compared to approximately 800 Miller units when the wild-type *UPC2A* gene was present. Loss of *UPC2A* lowered expression to approximately 80 Miller units. All *C. glabrata* isolates carrying the *CYC1-lacZ* control plasmid lacking an upstream activation sequence showed minimal levels of β-galactosidase activity. These data are consistent with this reporter system faithfully detecting the status of the *UPC2A* gene.

Next, we transformed these two plasmids into strains containing the various *ERG11* alleles. Appropriate transformants were grown to mid-log phase and enzyme levels determined (Figure 4B). The highest level of β-galactosidase activity was produced in the presence of the DM *ERG11* allele, consistent with this strain triggering the highest level of Upc2A activity. Both single mutant *ERG11* alleles induced smaller but significant increases in expression of the 5X SRE-*CYC1-lacZ* plasmid compared to the wild-type *ERG11*-containing strain. Only minimal levels of β-galactosidase activity were produced in strains containing the *CYC1-lacZ* reporter plasmid lacking any SREs as upstream control elements. These data support the view that the DM *ERG11* allele caused the largest increase in Upc2A function, likely through production of the most compromised form of the Erg11 enzyme, but each single mutant had a less dramatic but still detectable impact on Upc2A. To directly assess Upc2A function, we carried out single gene chromatin immunoprecipitation analysis of Upc2A binding to target promoters

Previous work from several labs including ours (20, 25) has provided evidence that ergosterol limitation increased Upc2A DNA-binding to nuclear target genes. We also found that these Upc2A target genes included the *PDR1* and *CDR1* genes (19), linking control of ergosterol biosynthesis with expression of genes involved in drug resistance. To determine if the presence of the fluconazole-resistant, DM allele of *ERG11* caused increased levels of Upc2A DNA binding, chromatin immunoprecipitation was used to detect binding of this factor to SREs present in the *ERG11*, *CDR1* and *PDR1* promoters. Non-specific binding was also evaluated by examining Upc2A enrichment on the HO promoter and *ERG11* coding sequence. Isogenic wild-type and DM *ERG11* strains were grown to mid-log phase and Upc2A DNA-binding analyzed by ChIP using anti-Upc2A or a non-specific control polyclonal antiserum (Figure 4C).

Upc2A binding to the *ERG11* SRE was strongly induced when cells contained the DM *ERG11* allele compared to the wild-type gene. Upc2A association with the *PDR1* SRE was also induced under these conditions although binding to *CDR1* did not change. Control ChIP reactions confirmed that the Upc2A binding was antibody-specific and showed no enrichment with either the *HO* gene or the *ERG11* coding sequence. The increased Upc2A-dependent transcriptional activation and DNA-binding indicate that Upc2A function was increased in strains containing the DM *ERG11* allele compared to the wild-type gene.

### Phenotypic comparison of *PDR1* GOF and DM *ERG11* mutations indicate mutant Pdr1 factors drive higher level azole resistance

Since the DM *ERG11* mutant was capable of inducing strong fluconazole resistance, we wanted to compare its behavior to that seen in the presence of *PDR1* GOF alleles. These GOF mutants of *PDR1* represent the vast majority of azole resistant mutants found in *C. glabrata* isolates (26–28). An isogenic series of strains was produced that produced wild-type or DM mutant forms of Erg11 containing the following alleles of *PDR1*: *pdr1Δ*, wild-type, D1082G or R376W. These strains were grown to mid-log phase and tested for their relative susceptibility to 30 μg/ml fluconazole in a serial dilution plate assay (Figure 5A).

**Figure 5.**
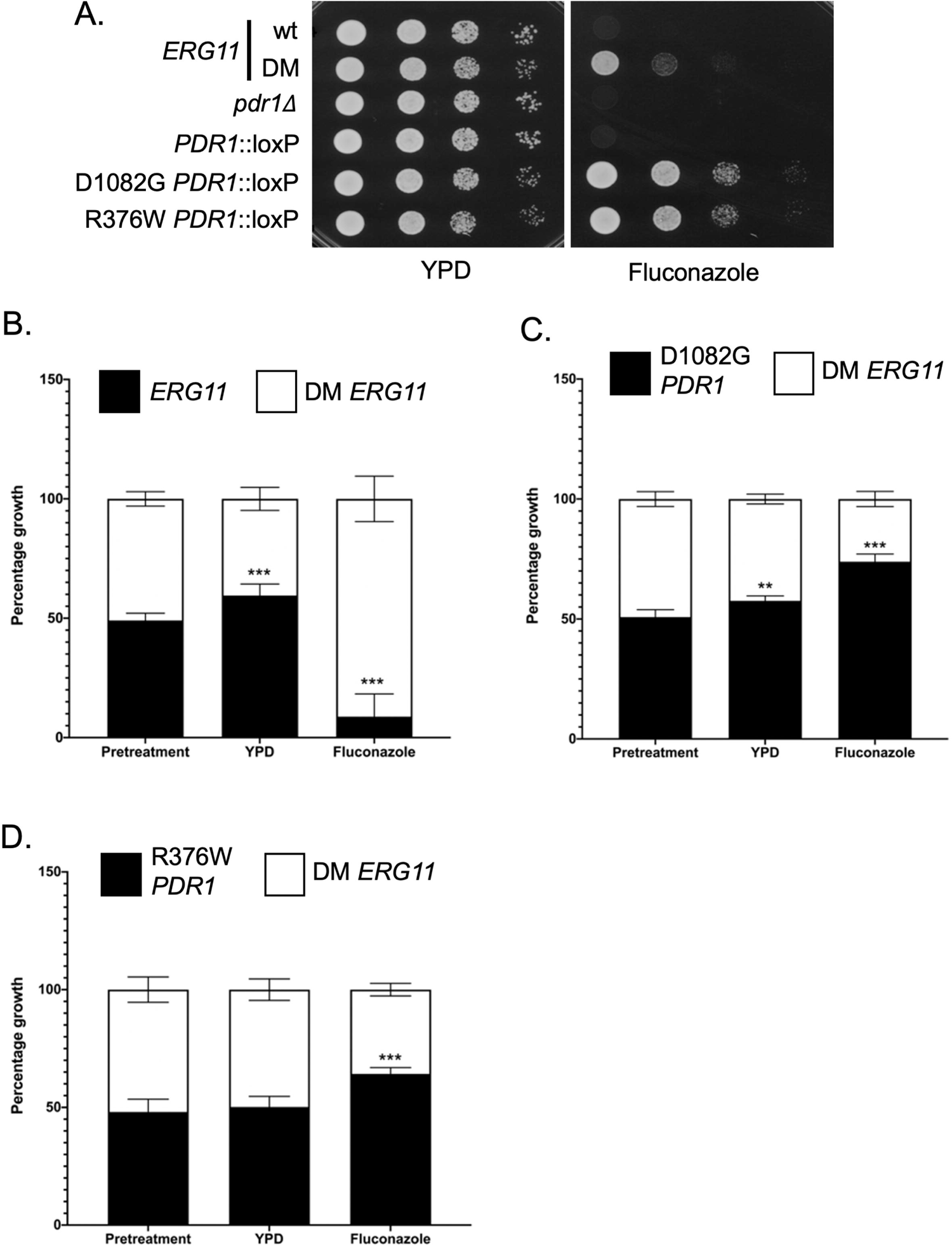
Growth phenotypic comparison of DM *ERG11* with gain-of-function (GOF) alleles of *PDR1*. A. Isogenic mid-log phase *C. glabrata* strains containing *ERG11* wild-type or), wild-type Δ*PDR1* (*PDR1*::loxP), or two GOF alleles of *PDR1* D1082G and R376W were serially diluted and spotted on YPD agar plates supplemented with or without fluconazole (20 μg/ml). Plates were incubated at 30°C and photographed. B. Fitness comparisons between the DM *ERG11* strain compared with wild-type (B), the GOF D1082G (C) and R376W (D) *PDR1* mutant strains. Equal number of mid-log phase cells from each strain were mixed together (Pretreatment). Cultures were then allowed to grow in the absence (YPD) or presence of azole drug (Fluconazole) (20 μg/ml). At the end of incubation, samples of each culture were plated and final distribution of each strain determined by plating. Deviations from the starting 50:50 mix of strains would indicate a relative fitness advantage for one strain versus the other.

Both GOF forms of *PDR1* lowered fluconazole susceptibility more than the DM *ERG11* strain. To probe the relative effects of mutations in either *PDR1* or *ERG11*, we carried out a competitive growth assay in which equal numbers of cells containing mutations of interest were mixed together. These mixed cultures were then allowed to grow in the presence or absence of fluconazole challenge and the relative contribution of each mutant strain to the final population determined. This competitive growth assay allows subtle differences to be more readily detected than on a plate-based assay.

We prepared mixed cultures of isogenic wild-type and DM *ERG11* cells (Figure 5B) and allowed these cultures to grow under untreated conditions (YPD) or in YPD medium containing fluconazole to late log phase. Growth in rich medium led to an increase in the population of the wild-type (60% final) compared to the DM *ERG11* strain (40% final). This analysis indicated that the DM *ERG11* had a growth defect compared to wild-type cells in the absence of fluconazole but this mutant allele conferred a significant growth advantage in the presence of this drug.

The relative growth of the DM *ERG11* strain was then compared to each of the *PDR1* GOF mutant strains (Figures 5C and 5D). In both cases, the GOF *PDR1* mutant outcompeted the DM *ERG11* strain in the presence of fluconazole (D1082G-74%: DM-26%; R376W-65%: DM-35%) while the only difference in growth in the absence of drug was seen in comparison of the D1082G *PDR1* strain compared to the DM *ERG11*: 58% to 42%, respectively. Both GOF *PDR1* mutants were more effective in conferring fluconazole resistance while growing at least as well (R376W) if not better (D1082G) than the DM *ERG11* strain. While we are only testing a single mutant form of *ERG11* here, growth differences associated with azole-resistant forms of Erg11 may help explain the predominance of GOF mutants of *PDR1* in clinical azole-resistant isolates of *C. glabrata*.

### Increased activation of *PDR1* and *CDR1* transcription seen in presence of GOF *PDR1* mutants compared to DM *ERG11*

The phenotypic data above indicated that both GOF *PDR1* strains produced a stronger effect on fluconazole resistance than the presence of the DM *ERG11* allele. To probe the basis of this phenotypic difference, we examined transcription of *CDR1*, *PDR1* and *ERG11* by RT-qPCR. Strains containing wild-type, D1082G or R376W forms of *PDR1* or the DM *ERG11* were grown to mid-log phase in the absence or presence of fluconazole. Levels of mRNA from these three genes were assayed as before.

*CDR1* mRNA levels were strongly elevated in the presence of both GOF *PDR1* mutations, well above those induced by the presence of the DM *ERG11* mutation (Figure 6A). This increase was seen both in the absence as well as the presence of fluconazole. *PDR1* mRNA levels were significantly higher in the presence of the two GOF alleles than in the DM *ERG11* but this difference was restricted to the absence of fluconazole. Drug treatment elevated both wild-type and DM *ERG11* strains to levels similar to those of the GOF mutants. Finally, *ERG11* mRNA levels were higher in the DM *ERG11* strain than in the other three strains in the absence of fluconazole. Strikingly, fluconazole exposure induced *ERG11* mRNA to higher levels in the strains containing wild-type *PDR1* but was less effective in the two GOF *PDR1*-containing strains. These data suggest that the increased *PDR1*/*CDR1* expression seen in the GOF *PDR1* strains may reduce the level of fluconazole stress (and associated *ERG11* induction) seen. The changes seen in mRNA levels were well-correlated with western blot analyses for each protein (Figures 6B and 6C). Upc2A protein levels were constant under all conditions tested, consistent with changes due to this factor being caused by alterations in Upc2A function rather than its expression.

**Figure 6.**
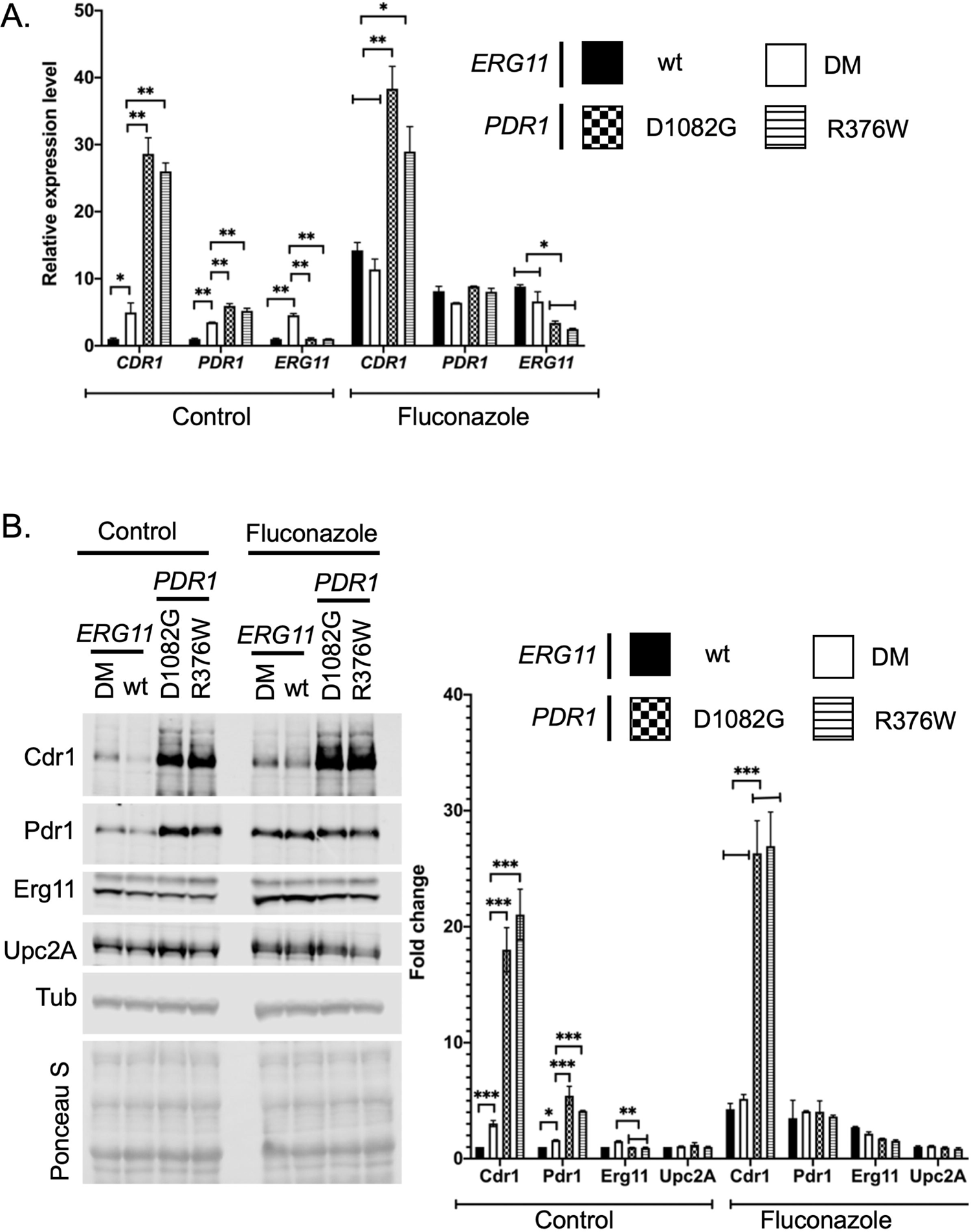
Relative gene expression responses to the DM *ERG11* or two different GOF *PDR1* strains. A. Strains varying at the *ERG11* or *PDR1* loci as indicated were grown in the presence or absence of fluconazole as in Figure 3 and processed for RT-qPCR analyses of the indicated mRNAs. B. The strains above were analyzed by western blotting as in Figure 3 using antisera for Cdr1, Pdr1, Erg11, Upc2A or Tubulin (Tub). The membrane was stained after transfer with Ponceau S as above. Quantitation of 4 western experiments is provided on the right hand side of the figure.

### Reduced levels of Upc2A exacerbate the growth defect of the DM *ERG11* strain

Our analysis of the cellular response to the presence of the DM *ERG11* mutation demonstrated the involvement of Upc2A as activity of this factor was induced in this genetic background compared to wild-type cells. We introduced the highly repressible *MET3* promoter (29) in place of the chromosomal *UPC2A* cognate region in order to test the effect of changing Upc2A production via repression in the presence of exogenous methionine. This *MET3-UPC2A* fusion gene was generated in both wild-type and DM *ERG11* strains. Appropriate transformants were placed on media lacking (SC) or containing excess methionine (SC+methionine) and allowed to grow. Two independent isolates of each *MET3-UPC2A* transformant were tested to evaluate strain variability in these assays.

Addition of methionine to strains containing the *MET3-UPC2A* fusion gene in addition to the DM *ERG11* allele caused the growth of this strain to be further reduced when compared to the presence of the wild-type *UPC2A* gene (Figure 7A, right hand panel). This result suggests that reduction of Upc2A levels (by repression of the *MET3* promoter with methionine) caused a growth defect in the presence of the DM *ERG11* mutation. When these *UPC2A* and *MET3-UPC2A* strains containing the DM *ERG11* mutation were grown in the absence of exogenous methionine, their growth was similar (Figure 7A left hand panel).

**Figure 7.**
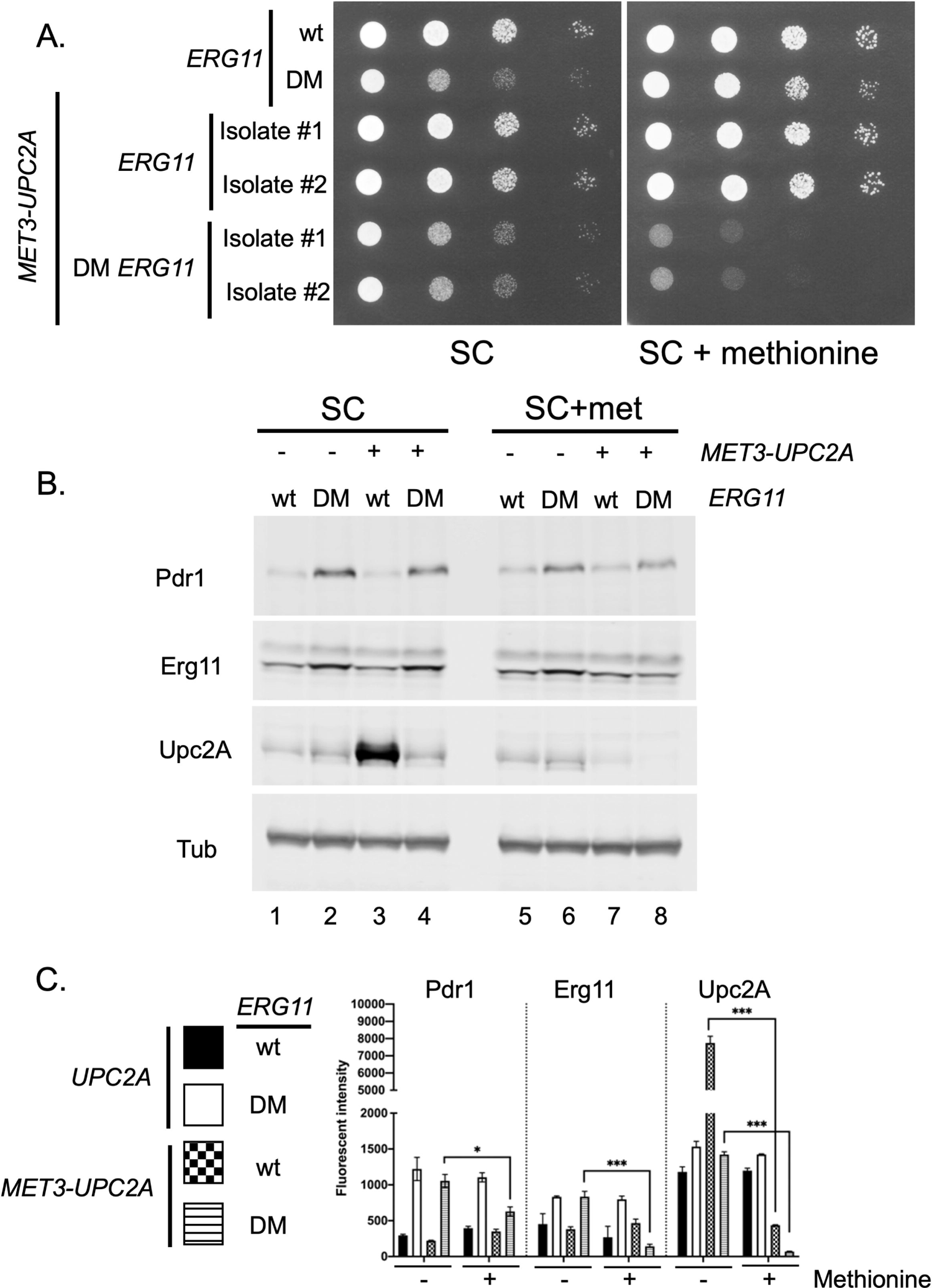
Depletion of Upc2A enhances growth defect of a DM *ERG11* strain. A. The *ERG11* promoter was replaced with the methionine-repressible *MET3* promoter in wild-type and DM *ERG11* cells. Two different isolates of each *MET3* promoter replacement were grown to mid-log phase, along with the isogenic wild-type and DM *ERG11* strains. These cells were serially diluted onto plates containing either synthetic complete (SC) medium or the same medium containing an excess of methionine (SC + methionine). Transformants were grown at 30°C and the plates photographed. B. The indicated strains were grown to mid-log phase in the presence or absence of methionine and whole cell protein extracts prepared. These extracts were analyzed as before using the indicated antisera. C. Quantitation of protein levels in multiple western blots is shown. Significant changes are indicated with asterisks.

To determine the levels of Upc2A protein in these different strains, we prepared whole cell protein extracts from these strains. These extracts were analyzed by western blotting using the anti-Upc2A antiserum. The *MET3* promoter activity is highest in SC medium (29) and we will consider the data from these media conditions first (Figure 7B, left hand panel). The *MET3* promoter drove much higher levels of Upc2A protein than were produced from the native *UPC2A* promoter in the presence of wild-type *ERG11* (compare lanes 1 and 3). The elevated expression of Upc2A seen in wild-type cells containing *MET3-UPC2A* did not lead to a corresponding increase in Erg11 expression (lanes 1 and 3). Interestingly, Upc2A levels were dramatically reduced when the *MET3-UPC2A* fusion gene was present in the DM *ERG11* strain, albeit to levels similar to those seen when Upc2A was produced by the native *UPC2A* gene (compare lanes 2 and 4). The presence of the *MET3-UPC2A* allele did not affect the observed induction of Erg11 in the presence of the DM *ERG11* allele (compare lanes 2 and 4), potentially due to the surprisingly low levels of Upc2A driven by the *MET3-UPC2A* allele when the DM *ERG11* mutant was present. The lack of any detectable Upc2A expression differences in the two DM *ERG11* strains (varying due to their different *UPC2A* promoters) also correlated well with the equivalent ability of these strains to grow on SC medium (Figure 7A above).

When methionine was added to repress *MET3-UPC2A* transcription, the levels of the corresponding Upc2A protein were strongly reduced as expected (Figure 7B, right hand panel). However, the reduction of Upc2A upon methionine repression led to different levels of this transcription factor in a manner dependent on the *ERG11* allele present in the cell. Methionine repression of the *MET3-UPC2A* in the DM *ERG11* strain blocked the production of a detectable level of Upc2A while this same treatment of cells carrying wild-type *ERG11* still produced readily detectable Upc2A (compare lanes 7 and 8). The extremely low level Upc2A production seen in the methionine-repressed *MET3-UPC2A* with DM *ERG11* may explain the methionine-dependent growth defect seen in this strain (see Figure 7A). These low levels of Upc2A also blocked the normal increase in Erg11 and Pdr1 caused by the DM *ERG11* allele (compare lanes 6 and 8), suggesting that both Erg11 and Pdr1 expression were, in part, driven by Upc2A function in this mutant background.

We carried out this same analysis but using a *MET3-PDR1* gene (Supplemental Figure 2). In the absence of fluconazole, there was no significant effect of the *MET3-PDR1* fusion gene on growth of either the wild-type or DM *ERG11* strain irrespective of the presence of methionine (Supplemental Figure 2A). The addition of fluconazole revealed the strong increase in resistance shown by the DM *ERG11* strains in the presence of either the wild-type or *MET3*-driven *PDR1* gene, in the absence of methionine (Supplemental Figure 2A, top panels). The addition of methionine to these media to repress *MET3-PDR1* (Supplemental Figure 2A, bottom panels) caused a loss of fluconazole resistance at the highest concentration tested. These data illustrate the critical contribution made by Pdr1 to the fluconazole resistance phenotype of the DM *ERG11* strain.

Western blot analyses were carried out to determine the effect of the *MET3-PDR1* gene on expression of Pdr1 itself, Cdr1 and Erg11 (Supplemental Figure 2B). Steady-state levels of Pdr1 were much higher in the presence of the wild-type *ERG11* gene than in the DM *ERG11*-containing strain (compare lanes 1 and 3). The expression of Pdr1 in these two strains was inversely correlated with expression of the Pdr1 target gene *CDR1* as evidenced by the high Cdr1 levels in the DM *ERG11* strain compared to the wild-type strain (compare lanes 1 and 3). This finding suggests that Pdr1 is activated in the presence of the DM *ERG11* allele as its steady-state expression is lower yet Cdr1 expression is higher. Previously, we used the *MET3* promoter to produce either wild-type or GOF Pdr1 proteins (30). The GOF proteins were found to be less stable than the corresponding wild-type factor. Production of this hypomorphic form of Erg11, in the absence of any azole drug, was sufficient to strongly induce Pdr1 activity and Cdr1 synthesis.

## Discussion

One of the nearly inevitable outcomes from the extensive use of fluconazole as an antifungal drug has been the development of resistance. Mutant forms of *ERG11* have been found in every *Candida* species with the exception of *C. glabrata* and *C. krusei* (31). Detailed analyses of the *C. albicans* Erg11 enzyme have clearly implicated structural changes within azole-resistant mutant forms of this enzyme leading to reduced azole susceptibility (reviewed in (32)). Azole resistance caused by mutational alteration within Erg11 is most parsimoniously interpreted in this manner.

Our data provide a second means of *ERG11* mutants impacting azole resistance via a secondary effect on expression of Pdr1/*CDR1* in *C. glabrata*. While it is clear that the mutants we have characterized here have effects in addition to activation of Pdr1, genetic depletion of Pdr1 caused an approximate 5-fold reduction in fluconazole resistance in the DM *ERG11* strain (Supplemental Figure 2). This type of a secondary effect can have major implications in the observed azole resistance of a given *ERG11* mutant background and is an important consideration in evaluation of this phenotype.

Fluconazole resistant clinical isolates of *C. albicans* are readily recovered with lesions in the Ca*ERG11* gene but azole resistant isolates with mutations in this locus have rarely been reported in *C. glabrata* (33). We constructed *C. glabrata ERG11* mutations based on cognate changes in the *C. albicans* gene. We selected positions Y141 and S410 in the *C. glabrata* enzyme as the analogous positions in the *C. albicans* Erg11 could be found mutated either alone or together in azole-resistant clinical isolates (9). The *C. glabrata* equivalent mutations caused azole resistance as expected but the DM allele also exhibited a growth defect, even on rich YPD medium, which was not reported for the *C. albicans* mutant. This growth defect, at least for the DM *ERG11* mutant, may be part of the reason that mutations in *ERG11* are not found in *C. glabrata* isolates. Confirmation of this suggestion will require more detailed study of the corresponding mutations in *C. albicans* as well as analyses of additional *ERG11* mutants in *C. glabrata*.

The DM *ERG11* allele was also found to cause several other drug phenotypes. The increased resistance to caspofungin was not expected and may involve induction of the *FKS2* gene (Figure 2D). We have preliminary evidence that Upc2A may act to control expression of β-glucan synthase (20) but have not yet linked this to *FKS2*. Crz1 is a well-established transcriptional activator of *FKS* gene expression (34, 35) and we provide evidence that the CRZ1 gene must be intact for the DM *ERG11* allele to induce caspofungin resistance (Figure 2E).

The DM *ERG11* seems to be acting as a reduced function or hypomorphic allele of this gene. This was evidenced by the reduced level of ergosterol accumulation in the DM mutant background and the increased amphotericin B resistance. We suggest that this reduction in function is critical for the observed induction of the *ERG* pathway genes as well as the Pdr1/*CDR1* regulatory circuit. Previous work from our lab showed that genetically depleting Erg11 by repression of two different artificial promoters driving *ERG11* caused the same changes in gene regulation (19). The behavior of the DM *ERG11* mutant strain provides further support for our hypothesis that a physiological link exists between the ergosterol biosynthetic pathway and Pdr1 (18).

Comparison of the DM *ERG11* allele with each of the two single mutant alleles suggests that there is a synthetic interaction between the Y141H and S410F mutations. In neither of the two single mutants was evidence obtained that endogenous downstream gene expression was being induced in the presence of these altered enzymes. However, when both of these lesions were present in the same protein, a range of downstream responses were triggered. These include *FKS2* induction and Pdr1 activation along with Upc2A. This was not observed with the single mutants. We suggest that the single mutants seem to cause increased fluconazole resistance primarily through their action on activity of the altered enzyme. The very sensitive 5X SRE-*CYC1-lacZ* reporter plasmid was modestly induced in the presence of each single mutant Erg11 (Figure 4B) but their effect was small compared to the induction caused by the DM Erg11 protein. The combination of the two single mutations led to production of a more defective Erg11 protein that in turn caused intracellular responses such as activation of Upc2A and Pdr1.

The activation of Upc2A-dependent transcription is a central feature of the cellular response to the presence of the DM *ERG11* allele. We produced a conditional *MET3-UPC2A* strain that contained the DM *ERG11* mutation to avoid any problematic interactions caused by introduction of a *upc2AΔ* null allele into this strain. The essential contribution of Upc2A to the maintenance of the DM *ERG11*-containing strain could be shown by the severe growth defect caused by methionine repression of this *UPC2A* allele (Figure 7A). We suggest that the presence of the DM *ERG11* allele led to induction of Upc2A function which in turn is linked to an increased level of degradation of this active transcription factor. This suggestion is supported by the inhibition of *MET3-UPC2A* transcription by the addition of methionine leading to a drop in Upc2A levels below the limit of detection in the DM *ERG11* background (Figure 7B). This is roughly the equivalent of a direct protein stability assay as we block synthesis at the level of transcription while allowing degradation to proceed without intervention. The apparent instability of the wild-type Upc2A after activation by the presence of the DM Erg11 protein is very similar to the instability found when GOF forms of Pdr1 are compared to wild-type Pdr1 (17). These factors are evidently deleterious for the cell to maintain in their activated states unless their presence is crucial for a given stress response. We have previously observed this as a hyperactive mutant form of Pdr1 can be lethal (17). These data suggest that an important control point for Upc2A may lie at the level of protein stability.

One reason for constructing these different fluconazole resistant forms of Erg11 in *C. glabrata* was to see if their presence would influence fluconazole induction of Pdr1. It is possible that fluconazole resistant forms of Erg11 would no longer respond to challenge by this azole drug. However, that was not the case as all strains expressing fluconazole resistant *ERG11* allele still showed induction of the Pdr1/*CDR1* regulatory axis upon fluconazole exposure (Figure 3). The striking induction of *CDR1*, *PDR1* and *ERG11* expression when the DM *ERG11* allele is present with no fluconazole addition argues that these genes and the pathways they define are normally co-regulated, irrespective of the presence of drug.

The simplest interpretation for azole resistant *ERG11* mutations in multiple organisms has been that the altered enzymes may no longer be able to interact with azole drugs and this can explain their resistance phenotype. While this is certainly true for some mutants (such as the two single mutant alleles of *ERG11* analyzed here), there are likely to be more complex physiological responses to these mutant enzymes that can alter the resistance phenotype in *ERG11* mutant strains. Ergosterol is an essential constituent of the fungal cell membrane and limiting its biosynthesis triggers transcriptional correction in most fungi examined (5, 36, 37). In *C. glabrata*, there appears to be a threshold of ergosterol limitation that is not crossed by the single mutant forms of Erg11 but is exceeded by the DM Erg11 enzyme. Exceeding this threshold leads to activation of downstream pathways to deal with the detected ergosterol limitation.

Our finding of these deeper regulatory connections between ergosterol biosynthesis and transcription circuitry including but not restricted to factors directly involved in *ERG* gene control suggests that additional linkages of this type may be participating in azole resistance in other fungi. Azole-resistant forms of *Aspergillus fumigatus* are commonly associated with substitution mutations within the *cyp51A* gene (*ERG11* equivalent in *Aspergilli*) (37). Some of the most common azole-resistant alleles found in *A. fumigatus* are the result of linked changes in which a residue in the coding sequence is changed and a region of the promoter is duplicated (Reviewed in (38)). These linked changes act together to enhance azole resistance more than either alone (39), suggesting that alleles of this sort engage more than one mechanism to affect azole resistance. We have previously shown that the L98H form of Cyp51A in *A. fumigatus* produces lower levels of this enzyme than the wild-type yet is still resistant to voriconazole (40). Additionally, mutations in the HMGCoA reductase gene (*hmg1*) from *A. fumigatus* also causes a strong decrease in voriconazole susceptibility although *cyp51A* gene expression appeared to be unaffected (41). These observations suggest that other fungi may also employ additional transcriptional circuitry to impact azole resistance in response to alterations in ergosterol biosynthesis. Identifying and characterizing these other transcriptional contributors to azole resistance are important goals in the dissection of the molecular basis of azole resistance.

## Materials and Methods

### Strains and growth conditions

*C. glabrata* was grown in rich YPD medium (1% yeast extract, 2% peptone, 2% glucose) or under amino acid-selective conditions in complete supplemental medium (CSM) (Difco yeast nitrogen extract without amino acids, amino acid powder from Sunrise Science Products, 2% μg/ml nourseothricin (Jena Bioscience, Jena, Germany) and 2 mM methionine was used to select strains containing pBV65 vector derivatives (42). CSM media (2% glucose, 1mM estradiol) without methionine was used to recycle the selection cassette on pBV65. YPD solid agar supplemented with 1 mg/ml 5-fluoro-orotic acid was used to cure the pBV43 plasmid. All strains used in this study are listed in Table 1.

**Table 1.**
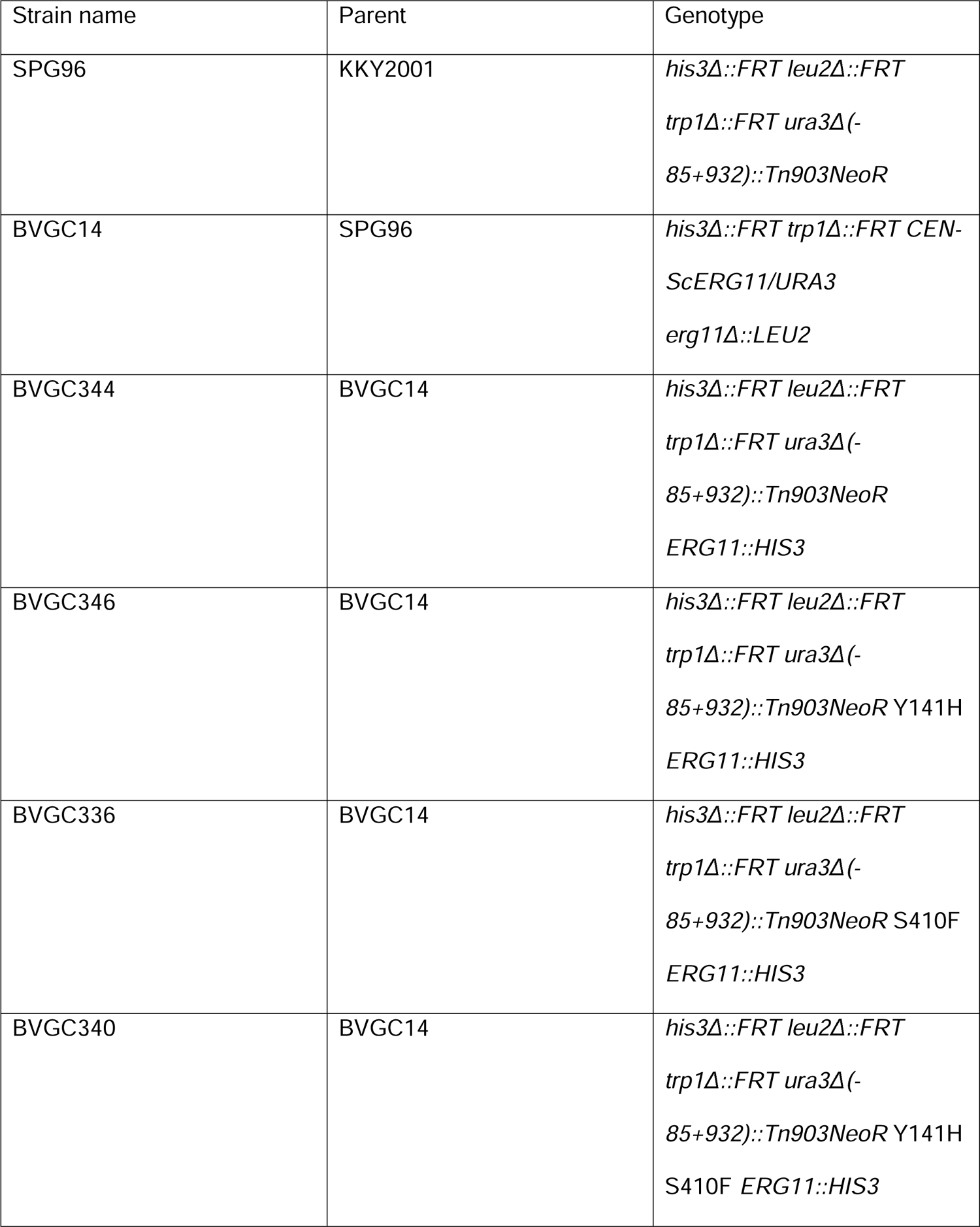
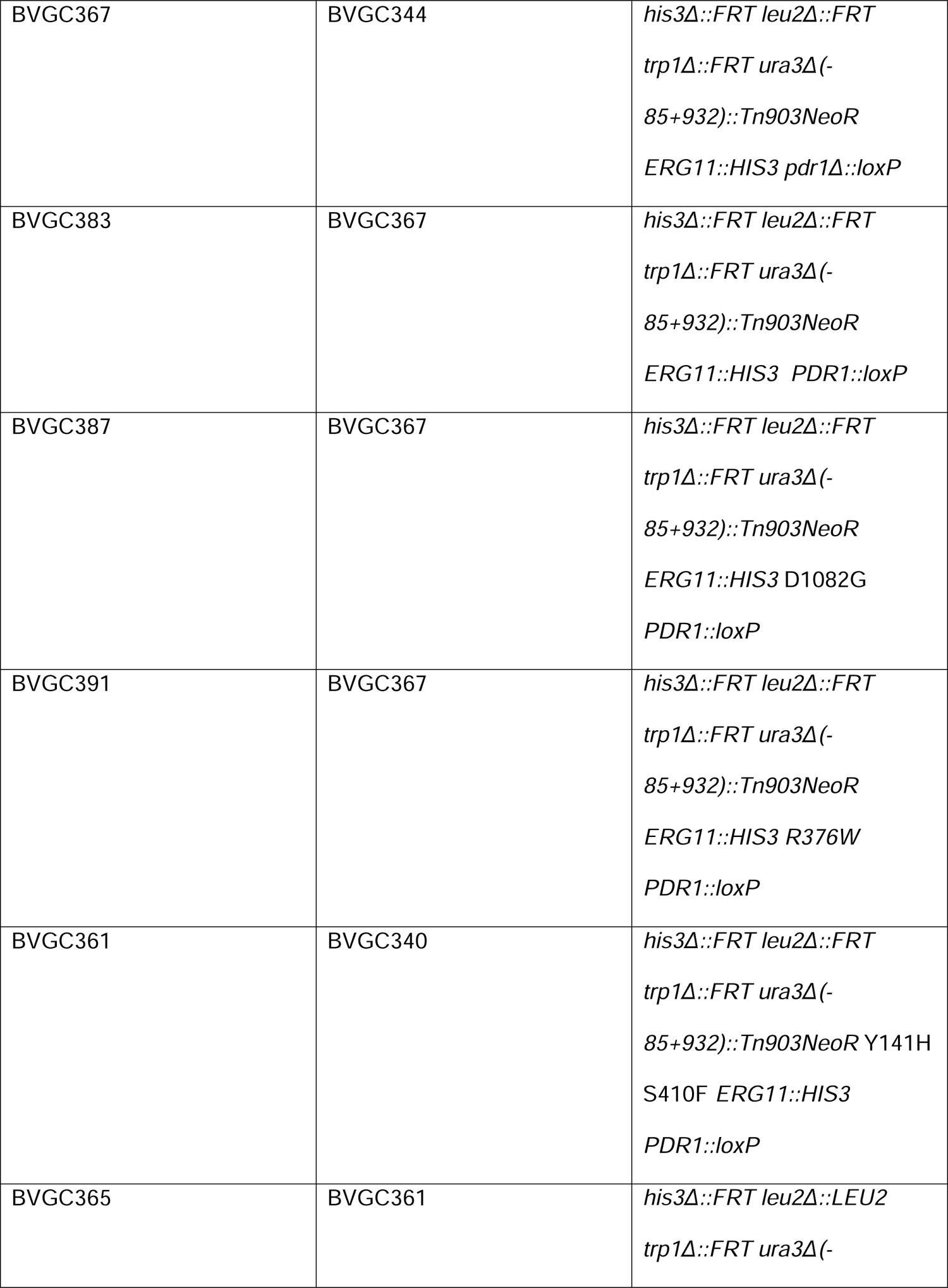
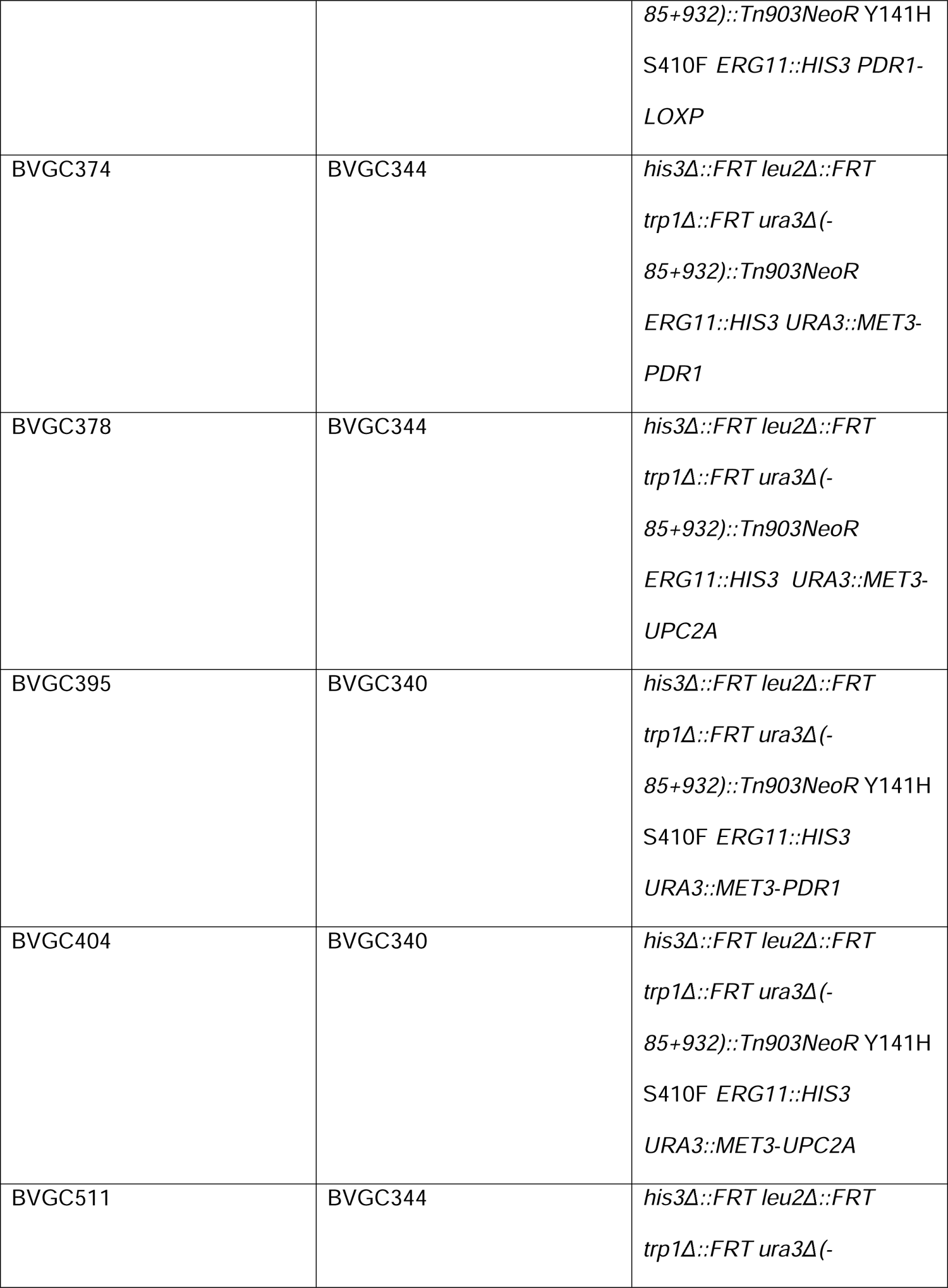
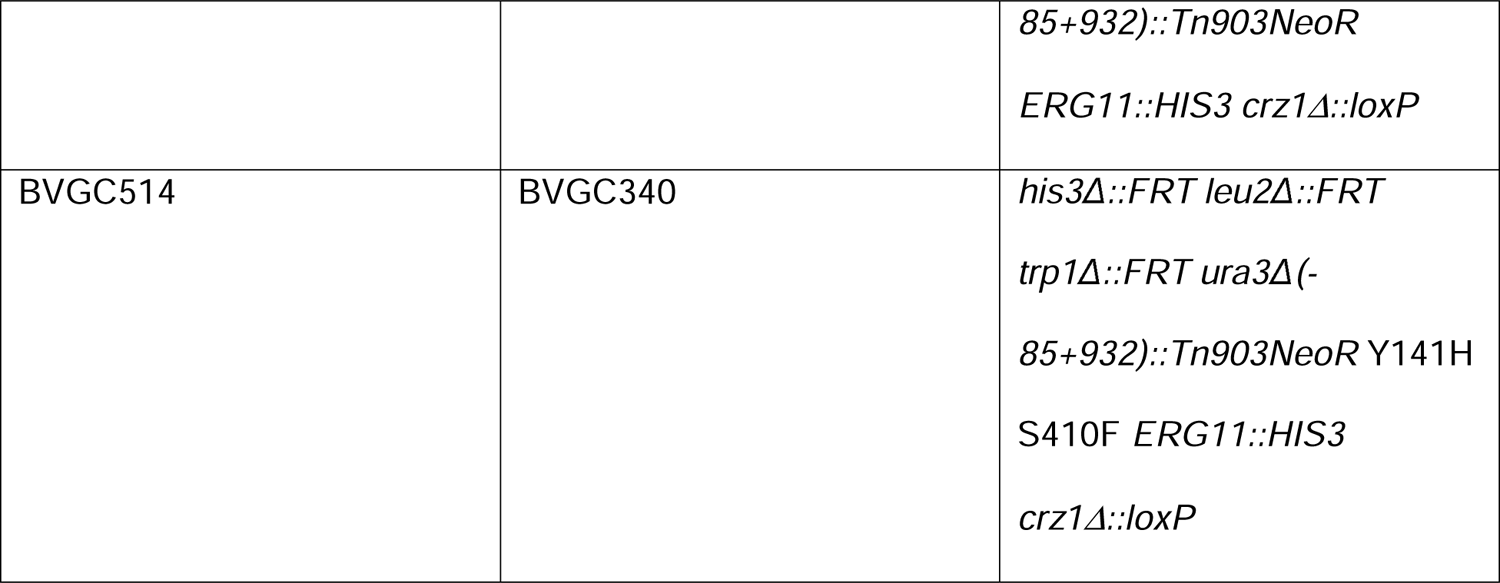
Strains used in this work.

| Strain name | Parent | Genotype |
| --- | --- | --- |
| SPG96 | KKY2001 | *his3Δ::FRT leu2Δ::FRT*<br>*trp1Δ::FRT ura3Δ(-*<br>*85+932)::Tn903NeoR* |
| BVGC14 | SPG96 | *his3Δ::FRT trp1Δ::FRT CEN-*<br>*ScERG11/URA3*<br>*erg11Δ::LEU2* |
| BVGC344 | BVGC14 | *his3Δ::FRT leu2Δ::FRT*<br>*trp1Δ::FRT ura3Δ(-*<br>*85+932)::Tn903NeoR*<br>*ERG11::HIS3* |
| BVGC346 | BVGC14 | *his3Δ::FRT leu2Δ::FRT*<br>*trp1Δ::FRT ura3Δ(-*<br>*85+932)::Tn903NeoR Y141H*<br>*ERG11::HIS3* |
| BVGC336 | BVGC14 | *his3Δ::FRT leu2Δ::FRT*<br>*trp1Δ::FRT ura3Δ(-*<br>*85+932)::Tn903NeoR S410F*<br>*ERG11::HIS3* |
| BVGC340 | BVGC14 | *his3Δ::FRT leu2Δ::FRT*<br>*trp1Δ::FRT ura3Δ(-*<br>*85+932)::Tn903NeoR Y141H*<br>*S410F ERG11::HIS3* |
| BVGC367 | BVGC344 | *his3Δ::FRT leu2Δ::FRT*<br>*trp1Δ::FRT ura3Δ(-*<br>*85+932)::Tn903NeoR*<br>*ERG11::HIS3 pdr1Δ::loxP* |
| BVGC383 | BVGC367 | *his3Δ::FRT leu2Δ::FRT*<br>*trp1Δ::FRT ura3Δ(-*<br>*85+932)::Tn903NeoR*<br>*ERG11::HIS3 PDR1::loxP* |
| BVGC387 | BVGC367 | *his3Δ::FRT leu2Δ::FRT*<br>*trp1Δ::FRT ura3Δ(-*<br>*85+932)::Tn903NeoR*<br>*ERG11::HIS3 D1082G*<br>*PDR1::loxP* |
| BVGC391 | BVGC367 | *his3Δ::FRT leu2Δ::FRT*<br>*trp1Δ::FRT ura3Δ(-*<br>*85+932)::Tn903NeoR*<br>*ERG11::HIS3 R376W*<br>*PDR1::loxP* |
| BVGC361 | BVGC340 | *his3Δ::FRT leu2Δ::FRT*<br>*trp1Δ::FRT ura3Δ(-*<br>*85+932)::Tn903NeoR Y141H*<br>*S410F ERG11::HIS3*<br>*PDR1::loxP* |
| BVGC365 | BVGC361 | *his3Δ::FRT leu2Δ::LEU2*<br>*trp1Δ::FRT ura3Δ(-* |
|  |  | *85+932)::Tn903NeoR Y141H*<br>*S410F ERG11::HIS3 PDR1-*<br>*LOXP* |
| BVGC374 | BVGC344 | *his3Δ::FRT leu2Δ::FRT*<br>*trp1Δ::FRT ura3Δ(-*<br>*85+932)::Tn903NeoR*<br>*ERG11::HIS3 URA3::MET3-*<br>*PDR1* |
| BVGC378 | BVGC344 | *his3Δ::FRT leu2Δ::FRT*<br>*trp1Δ::FRT ura3Δ(-*<br>*85+932)::Tn903NeoR*<br>*ERG11::HIS3 URA3::MET3-*<br>*UPC2A* |
| BVGC395 | BVGC340 | *his3Δ::FRT leu2Δ::FRT*<br>*trp1Δ::FRT ura3Δ(-*<br>*85+932)::Tn903NeoR Y141H*<br>*S410F ERG11::HIS3*<br>*URA3::MET3-PDR1* |
| BVGC404 | BVGC340 | *his3Δ::FRT leu2Δ::FRT*<br>*trp1Δ::FRT ura3Δ(-*<br>*85+932)::Tn903NeoR Y141H*<br>*S410F ERG11::HIS3*<br>*URA3::MET3-UPC2A* |
| BVGC511 | BVGC344 | *his3Δ::FRT leu2Δ::FRT*<br>*trp1Δ::FRT ura3Δ(-* |
|  |  | *85+932)::Tn903NeoR*<br>*ERG11::HIS3 crz1Δ::loxP* |
| BVGC514 | BVGC340 | *his3Δ::FRT leu2Δ::FRT*<br>*trp1Δ::FRT ura3Δ(-*<br>*85+932)::Tn903NeoR Y141H*<br>*S410F ERG11::HIS3*<br>*crz1Δ::loxP* |

### Plasmid construction and promoter mutagenesis

All constructs used for homologous recombination into the chromosome were constructed in the pUC19 vector (New England Biolabs, Ipswich, MA). PCR was used to amplify DNA fragments.

PCR products were run on 1.5 % agarose gel (RPI, Troy, NY), excised and purified with Purelink quick gel extraction and PCR cleaning combo kit (Invitrogen, Carlsbad, CA). Gibson assembly cloning (New England Biolabs) was routinely used to assemble fragments together into appropriate plasmid backbones. All isogenic deletion constructs were made by assembling the recyclable cassette from pBV65 and fragments from the immediate upstream and downstream regions of the target genes. Fragments of the target genes were amplified from CBS138 background. Eviction of the recyclable cassette left a single copy of *loxP* in place of the excised target gene coding region.

*PDR1* constructs were made by using Gibson assembly (43) to produce fragments corresponding to the wild-type, R376W or D1082G alleles of *PDR1* with a recyclable cassette located downstream from the natural stop codon. Eviction of the recyclable cassette in the integrated constructs left a single copy of *loxP* 250-300 base-pairs downstream of the target gene stop codons. R376W and D1082G forms of *PDR1* were amplified from pSK70 and pSK71 backgrounds (17).

*ERG11* wild-type and mutant integration constructs were made as above with the resulting fragments corresponding to the wild-type, Y141H, S410F or the Y141H S410F alleles of *ERG11* with the *his3MX6* (19) marker located 270 bp from the natural stop codon.

Conversion of strains to *LEU2* was done by PCR amplifying the *LEU2* coding region along with 500 base pairs immediately upstream and downstream from the CBS138 background. Linear DNA was then transformed into KKY2001 and the colonies were selected on CSM agar without leucine.

To generate the *S. cerevisiae ERG11* complementing plasmid (pBV43), the Sc*ERG11* promoter (800 bps upstream of AUG), Sc*ERG11* coding sequence, and Sc*ERG11* terminator (350 bps downstream of the stop codon) were PCR amplified from the an S288c wild-type strain (SEY6210). This fragment was then cloned into the pCU-MET3 vector (29), which was pre-digested with SacI and XhoI.

The Sterol Response Element (SRE) reporter construct (pBV382) was produced by cloning the E. coli *lacZ* gene as a PCR fragment from pSK80 (17). This fragment was then cloned into the pCL vector (29), which was pre-digested with SaII and XhoI. A Upc2A-responsive promoter cassette, containing the 5 concatemerized copies of the *ERG1* promoter SRE (−725 to −775) located downstream from the *ADH1* terminator and upstream of the *CYC1* minimal promoter, was generated by Genscript (Piscataway, NJ). The Upc2A responsive promoter was PCR amplified and cloned into the pCL-*lacZ* vector, digested with SacI and SalI. The control vector (pBV378) was constructed by cloning the *ADH1* terminator and the *CYC1* minimal promoter, PCR amplified from BVGC7 (19), into the pCL-*lacZ* vector, digested with SacI and SalI.

To generate the *MET3*-driven *UPC2A* and *PDR1* strains, the *MET3* promoter and the *URA3* gene were PCR amplified from the pUC-*MET3* plasmid. This *URA3-MET3* promoter cassette was placed immediately upstream of the target gene AUG codon by recombination into the gene of interest.

### *C. glabrata* transformation

Cell transformations were performed using a lithium acetate method. After being heat shocked, cells were either directly plated onto selective CSM agar plates (for auxotrophic complementation) or grown at 30°C at 200 rpm overnight (for nourseothricin selection).

Overnight cultures were then plated on YPD agar plates supplemented with 50 μ nourseothricin. Plates were incubated at 30°C for 24 to 48 hours. In case of chromosomal insertion, individual colonies were isolated and screened by PCR for correct insertion of the targeted construct.

### Construction of mutant *ERG11* strains

Since Erg11 is essential for *C. glabrata* aerobic growth (44), A *URA3* (*ScURA3*)-containing plasmid shuffling technique was used to construct *ERG11* mutant alleles in the SPG96 (*ura3*Δ) *C. glabrata* strain (17). An autonomous plasmid pBV43 marked with *ScURA3* and containing the wild-type Sc*ERG11* was first transformed into the SPG96 strain. Then *ERG11* was deleted in the chromosome by homologous recombination, leaving Sc*ERG11* on pBV43 functioning as the sole copy of the *ERG11* gene. Next, the wildtype *C. glabrata ERG11* and the 3 mutant alleles were individually reintroduced to the normal locus by homologous recombination with selection for His+ transformants. Finally, the pBV43 plasmid was cured with 1 mg/ml 5-FOA on a YPD agar plate to construct the *C. glabrata* strains, BVGC344 (*ERG11*), BVGC346 (Y141H *ERG11*), BVGC336 (S410F *ERG11*), and a strain containing an ERG11 gene with both of these mutations present (BVGC340: DM *ERG11*).

### Total sterol estimation

Cell total sterol was extracted and measured as previously described (19). In short, cell pellets were lysed in 25% alcoholic potassium hydroxide at 90°C for 2 detected by spectrophotometric scanning between the wavelengths of 240 nm and 300nm.

The presence of ergosterol in the extracted sample resulted in a four-peak curve with peaks located at approximately 262, 270, 281, and 290 Quantification of transcript levels by RT-qPCR.

Total RNA was extracted from cells by extraction using TRIzol (Invitrogen) and chloroform (Fisher Scientific, Hampton, NH) followed by purification with RNeasy minicolumns (Qiagen, μg total RNA was reverse-transcribed using an iScript cDNA synthesis kit (Bio-Rad, Des Plaines, IL). Assay of RNA via quantitative PCR (qPCR) was performed with iTaq universal SYBR green supermix (Bio-Rad). Target gene transcript levels were normalized to transcript levels of 18S rRNA.

### Spot test assay

Cells were grown in YPD medium to mid-log-phase. Cultures were then 10-fold serially diluted and spotted onto YPD agar plates containing different concentrations of fluconazole (LKT laboratories, St Paul, MN), caspofungin (Apexbio, Houston, TX) or Amphotericin B (Sigma). All agar plates were incubated at 30°C for 24 to 48 h before imaging was performed.

### β-galactosidase assay

Harvested cells were lysed with glass beads (Scientific Industries Inc) in breaking buffer (100 mM Tris pH8, 1 mM Dithiothreitol, and 20% Glycerol) at 4°C for 10 min. Lysate was collected β-galactosidase enzymatic reaction was carried out in Z-buffer (60 mM Na_2_HPO_4_, 40 mM NaH_2_PO_4_, 10 mM KCl, 1 mM MgSO_4_, 50 mM 2-Mercaptoethanol) with 650 µg/ml O-nitrophenyl-β-D-galactopyranoside (ONPG). Miller units were calculated based on the equation: (OD_420_ x 1.7) / (0.0045 x total protein concentration x used extract volume x time). Bradford assay (Bio-Rad) was used to measure the total protein concentration in the lysate.

### Chromatin immunoprecipitation (ChIP)

The detailed ChIP protocol was previously described (19). The sheared chromatin was incubated with rabbit polyclonal anti-Upc2A antibody (1:50 dilution) or rabbit IgG control antibody, NBP2-24891 (Novus Biologicals, Centennial, CO) for 2L together with 30 l of washed protein G Dynabeads (Life Technologies) overnight on a nutator Lμ at 4^°^ C. Washing, reversal of cross-links, and purification of DNA processed by the use of ChIP were performed as described in (19).

Real-time PCR was performed on ChIP purified DNA, under the following conditions: 1Lcycle of 95°C for 30 s followed by 40 cycles of 95°C for 15 s and 57°C for 30 s on a MyiQ2 Bio-Rad μl volume of the DNA processed by the use of ChIP (diluted 5-fold) or of input (diluted 20-fold) DNA was used in a reaction mixture with a 20 μM concentration of each primer. The percent input method was used to calculate the signal of enrichment of the promoter region for each gene. *ERG11*, *CDR1*, and *PDR1* promoters were analyzed with primers specifically targeting the *ERG11* promoter (−561 to −694 relative to the ATG as +1), *CDR1* promoter (−476 to −665), and *PDR1* promoter (−551 to −651) regions. A region of *ERG11*, located within the coding sequence (+939 to +1042) and the *HO* promoter (−585 to −751) region were used as negative controls.

### Competitive growth assay

The *LEU2* coding region along with its immediate 500 bps up- and downstream sequences were amplified from CBS138 genomic DNA. The product was then used to transform the *leu2Δ*::FRT allele to *LEU2* in DM *ERG11* background at the *LEU2* native locus to generate BVGC365 (DM *ERG11*/*LEU2)*. A mid-log growth culture [between O.D. of 1 and 2] was diluted to 0.2 O.D. in fresh YPD. Each culture to be tested: wild-type, R376W *PDR1* or D1082G *PDR1*, was mixed at a 1:1 ratio with DM *ERG11*/*LEU2*, and the mixed cultures were treated either with fluconazole μg/ml) or ethanol for 24 hours at 30°C. Cultures were collected, serially diluted, and plated on YPD and CSM media without leucine for colony forming unit analysis.

### Antibodies and western immunoblotting

Cells were lysed with lysis buffer (1.85 M NaOH, 7.5% 2-Mercaptoethanol). Proteins were precipitated with 50% Trichloroacetic acid and resuspended in Urea buffer (40 mM Tris pH8, 8.0 M Urea, 5% SDS, 1% 2-Mercaptoethanol). Cdr1, Pdr1, and Upc2A rabbit polyclonal antibodies were previously described (19). Anti-HA monoclonal antibody was purchased from Invitrogen.

Secondary antibodies were purchased from LI-COR Biosciences. Imaging was performed with Odyssey CLx Imaging System (LI-COR Biosciences) and analyzed by Image Studio Lite Software (LI-COR Biosciences). Detected target band fluorescence intensity was normalized against tubulin fluorescence intensity and compiled from two biological replicate experiments and two technical replicates in each experiment, giving four replicates in total. Erg11 anti-peptide rabbit polyclonal antibody was produced by GenScript (Piscataway, NJ). The peptide sequence was AKIYWEKRHPEQKY, and it covered the 520th to 533rd amino-acids in the Erg11 protein primary sequence. The peptide antibody was confirmed by western blotting comparing wild-type and a strain carrying an *ERG11*-3X HA allele. Anti-peptide antibody detected native Erg11 from wild-type cells and the increased molecular mass of the Erg11-3X HA protein. Dual color LI-COR secondary antibodies were used to overlap the band intensity of Erg11 protein detected with anti-HA mouse monoclonal antibody and Erg11 peptide rabbit polyclonal antibody

## Statistics

The Student T-test was used to assess the statistical significance of results of comparisons of samples. Paired conditions were used for comparisons of results from the same isolate obtained under different treatment conditions, while unpaired conditions were used for comparisons of results from isolates obtained under the same treatment conditions (*, P < 0.5; **, P < 0.01; ***, P < 0.001).

## Acknowledgements

This work was supported by NIH AI152494 to WSM. We thank Dr. Damian Krysan for useful suggestions and Drs. Tom Conway and Lucia Simonicova for critically reading this manuscript.

## Supplemental Figure Legends

**Supplemental Figure 1. Voriconazole and itraconazole resistance among *ERG11* mutations**. Mid-log phase *C. glabrata* strains containing the indicated *ERG11* alleles were serially diluted and spotted on YPD agar plates supplemented with various concentrations of voriconazole (A) or itraconazole (B).

**Supplemental Figure 2. Interaction of DM *ERG11* allele and Pdr1.** A. The *PDR1* promoter was replaced with the *MET3* promoter in strains containing either the wild-type or DM *ERG11* allele. Strains were grown to mid-log phase and then serial dilutions plated on SC medium lacking or containing methionine. Fluconazole was added to these plates where indicated. Plates were incubated at 30°C and then photographed. B. The strains above were grown to mid-log phase in SC medium lacking methionine, methionine was either omitted (-) or added (+) and cultures allowed to continue to grow for equal times. Whole cell protein extracts were then prepared and analyzed by western blotting using the indicated antibodies.

