## Supplementary figures and images for "Multiple mechanisms impact fluconazole resistance of mutant Erg11 proteins in Candida glabrata"

### Supplemental Figure 1

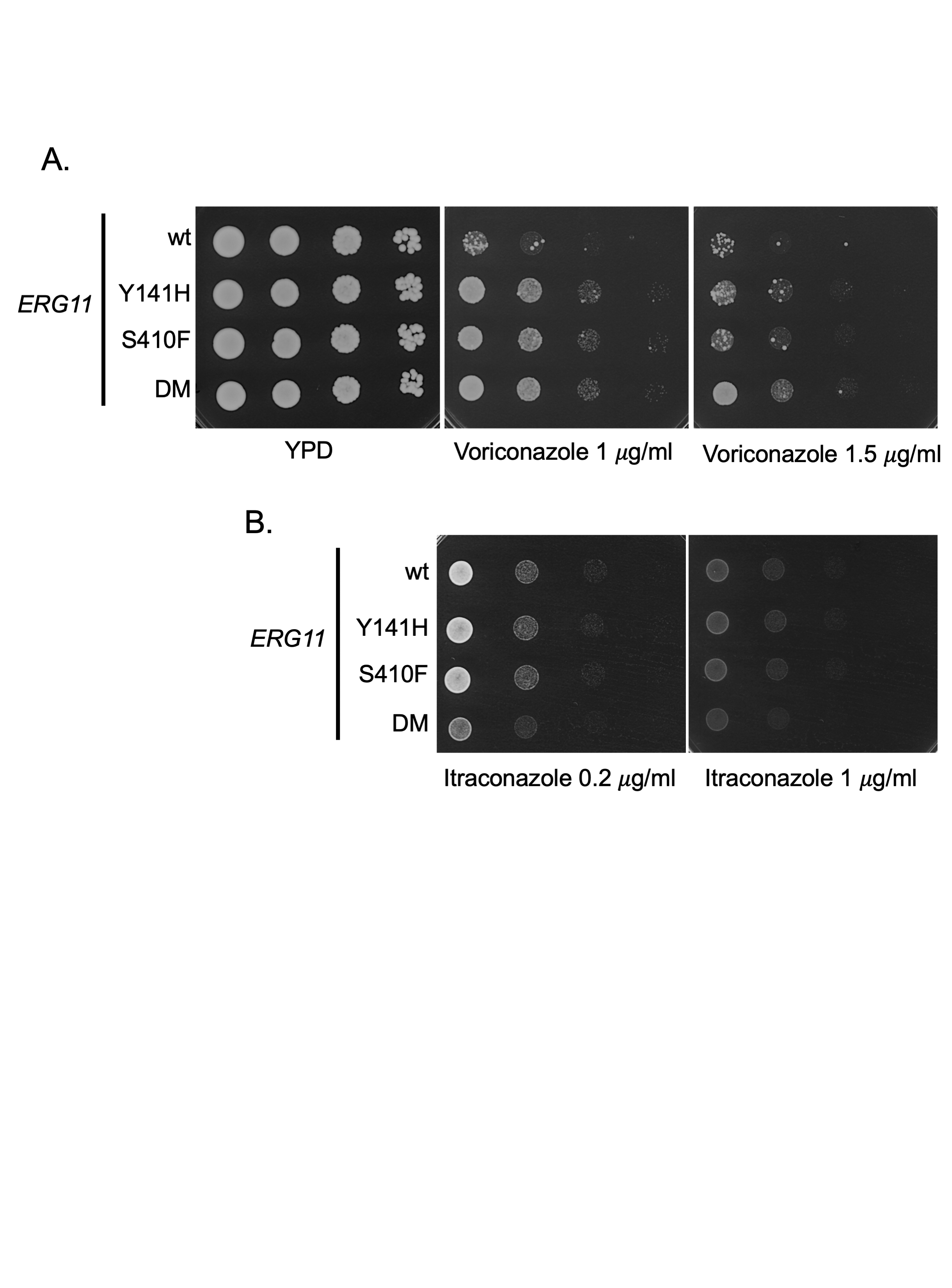

### Supplemental Figure 2

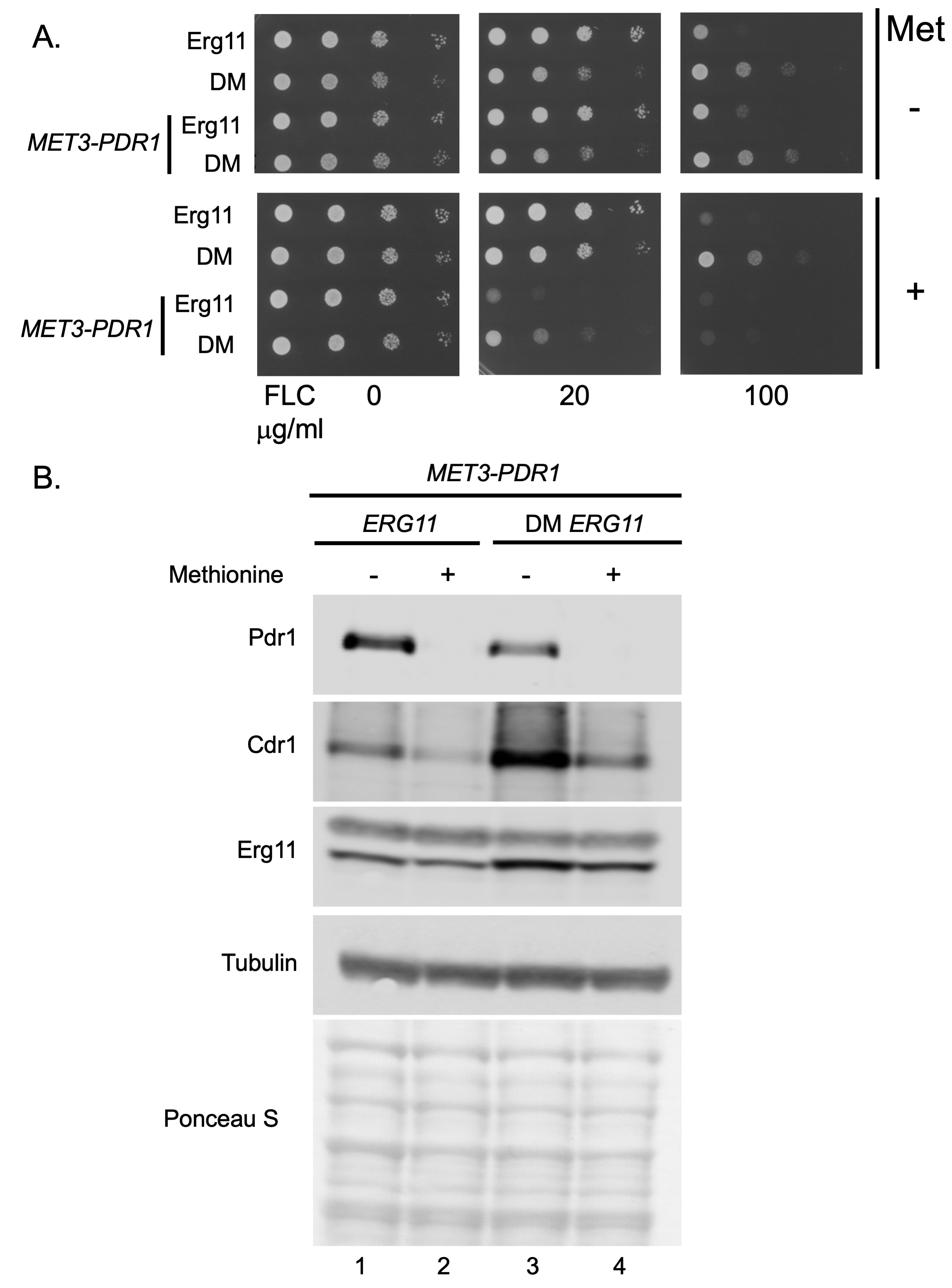
